# Differential reconfiguration of brain networks in children in response to standard versus rewarded go/no-go task demands

**DOI:** 10.1101/2024.10.15.618248

**Authors:** Mackenzie E Mitchell, Teague R Henry, Nicholas D Fogleman, Cleanthis Michael, Tehila Nugiel, Jessica R Cohen

## Abstract

Response inhibition and sustained attention are critical for higher-order cognition and rely upon specific patterns of functional brain network organization. This study investigated how functional brain networks reconfigure to execute these cognitive processes during a go/no-go task with and without the presence of rewards in 26 children between the ages of 8 and 12 years. First, we compared task performance between standard and rewarded versions of a go/no-go task. We found that the presence of rewards reduced commission error rate, a measure considered to indicate improved response inhibition. Tau, thought to index sustained attention, did not change across task conditions. Next, changes in functional brain network organization were assessed between the resting state, the standard go/no-go task, and the rewarded go/no-go task. Relative to the resting state, integration decreased and segregation increased during the standard go/no-go task. A further decrease in integration and increase in segregation was observed when rewards were introduced. These patterns of reconfiguration were present globally and across several key brain networks of interest, as well as in individual regions implicated in the processes of response inhibition, attention, and reward processing. These findings align with patterns of brain network organization found to support the cognitive strategy of sustained attention, rather than response inhibition, during go/no-go task performance and suggest that rewards enhance this organization. Overall, this study used large-scale brain network organization and a within-subjects multi-task design to examine different cognitive strategies and the influence of rewards on response inhibition and sustained attention in late childhood.

## Introduction

Although many foundational cognitive processes, including response inhibition and sustained attention, are present in a basic form in early childhood, they are less effective and consistent in childhood relative to adulthood (Levin et al. 1991; Tamm et al. 2002; Luna et al. 2004; Lorsbach and Reimer 2010; Bezdjian et al. 2014; Casey 2015; Luna et al. 2015; Lewis et al. 2017a; Lewis et al. 2017b). Both response inhibition and sustained attention have been shown to rely upon connectivity within and across distributed brain networks (Stevens et al. 2007; Marek et al. 2015; Chung et al. 2020; Thomson et al. 2022). Additionally, motivational supports, such as the presence of external rewards, enable improvements in both response inhibition and sustained attention in children (Kohls et al. 2009; Luciana and Collins 2012; Winter and Sheridan 2014; Demurie et al. 2016; Miyasaka and Nomura 2019; Burton et al. 2021; Sader et al. 2022). As such, understanding how intrinsic brain organization adapts to the cognitive demands of response inhibition and sustained attention, as well as how rewards elicit further changes in brain organization, will contribute to our understanding of the brain network mechanisms underlying these processes in children. To address this, we utilized standard and rewarded versions of a go/no-go fMRI task. Although go/no-go tasks are associated with an array of cognitive process, including working memory (Criaud and Boulinguez 2013) and motor execution (Simmonds et al. 2008; Wessel 2018), these tasks have primarily been used to probe response inhibition and sustained attention (Simmonds et al. 2008; Smith et al. 2013). The co-occurrence of response inhibition and sustained attention demands in go/no-go tasks make this an ideal medium for understanding how children engage in goal-driven behavior.

### Functional connectivity related to response inhibition and sustained attention in children and adolescents

Patterns of functional connectivity have been linked with response inhibition during go/no-go (Stevens et al. 2007; Barber et al. 2013; Cai et al. 2021) and related tasks (e.g., antisaccade, Stroop, and stop signal tasks; Mennes et al. 2012; Dwyer et al. 2014; Cai et al. 2019; Pan et al. 2020; Wang et al. 2021). Response inhibition, the ability to suppress a prepotent or automatic motor process, is operationalized in go/no-go tasks as accuracy on no-go trials (Stevens et al. 2007; Barber et al. 2013; Cai et al. 2021), on stop signal tasks as stop signal response time (Cai et al. 2019; Wang et al. 2021) or, conversely, stop signal delay (Mennes et al. 2012), or on interference tasks as performance decrement on incongruent trials (Dwyer et al. 2014; Pan et al. 2020). These studies have reported that stronger resting state (Mennes et al. 2012; Pan et al. 2020) and task-based (Dwyer et al. 2014; Cai et al. 2021) functional connectivity within and between regions of cognitive control-related networks (i.e., fronto-parietal and cingulo-opercular) is related to better response inhibition performance in children and adolescents. Additionally, stronger resting state (Mennes et al. 2012) and task-based (Stevens et al. 2007; Dwyer et al. 2014) anticorrelations between regions of the cognitive control-related networks and the default mode network have been found to be related to better response inhibition. Finally, it has been reported that stronger task-based functional connectivity between regions of cognitive control networks and subcortical regions, including the striatum (Wang et al. 2021) and subthalamic nucleus (Cai et al. 2019), relates to better response inhibition.

Studies have also used behavioral performance on go/no-go tasks and similar paradigms (e.g., continuous performance and stop signal tasks) to probe sustained attention in children and adolescents (Lewis et al. 2017a; Lewis et al. 2017b). Sustained attention is the ability to maintain goal-directed focus over a period of time and is often operationalized during cognitive tasks as low response time variability (Lewis et al. 2017a; Thomson et al. 2022). Functional connectivity studies provide evidence that reduced resting state (Barber et al. 2015; Thomson et al. 2022) and stop signal task (O’Halloran et al. 2018) functional connectivity strength between widely distributed brain networks is related to increased sustained attention. Brain networks implicated include those involved in cognitive control, attention, motor, and visual processing, as well as the default mode network (Barber et al. 2015; O’Halloran et al. 2018; Thomson et al. 2022). For example, it has been reported that reduced functional connectivity between the fronto-parietal and the default mode, dorsal attention, and limbic networks, as well as between the salience and the somatomotor networks, is related to reduced response time variability (Thomson et al. 2022). Additionally, it has been found that stronger within-network functional connectivity of the visual and auditory networks during passive movie watching is related to increased sustained attention as assessed by the Early Childhood Attention Battery (Breckenridge et al. 2013) in children aged 4-7 years (Rohr et al. 2018).

Together, studies investigating functional connectivity underlying response inhibition and sustained attention in children and adolescents implicate connections within and between several distinct brain networks. Stronger functional connectivity between cognitive control networks and weaker functional connectivity between the default mode and the cognitive control networks is related to better response inhibition. In contrast, reduced functional connectivity between widely distributed networks is associated with better sustained attention. To gain more insight into brain network topology underlying response inhibition and sustained attention, network-based metrics designed to probe large-scale, complex patterns of connectivity within and between several networks simultaneously have been utilized.

### Network organization related to response inhibition and sustained attention

Network-based studies have demonstrated that metrics describing network integration (i.e., connectivity across distinct brain networks) and network segregation (i.e., connectivity within distinct brain networks) are relevant to cognition (Sporns 2013; Cohen and D’Esposito 2016; Shine and Poldrack 2018). For example, participation coefficient is a measure commonly utilized that quantifies how broadly distributed a node or a network’s connections are to other brain networks. In adults, increased participation coefficient of the dorsal and ventral attention networks during the resting state is related to better response inhibition on a stop signal task (Hsu et al. 2020). Further, increased participation coefficient of the cingulo-opercular network during the resting state is related to better response inhibition during an antisaccade task in adolescents (Marek et al. 2015). As related to sustained attention, there is evidence from network-based studies in adults that periods of increased sustained attention are characterized by reduced integration (i.e., participation coefficient) and increased segregation (i.e., within module degree) of the cingulo-opercular, salience, dorsal attention, and visual networks, with the opposite pattern observed in auditory and somatomotor networks (Zuberer et al. 2021). Additionally, increased global segregation of brain networks (i.e., modularity) relates to more accurate stimulus detection during sustained perceptual vigilance tasks (Sadaghiani et al. 2015; Kucyi et al. 2018). Increased segregation has also been linked to tonic alertness (Sadaghiani & D’Esposito, 2015; Nie et al., 2022), a process inherent to sustained attention (Oken et al. 2006).

Thus, findings from studies reporting network-based metrics support and extend the findings from studies examining functional connectivity strength, in observing generally that higher network integration, or between-network connectivity, is important for response inhibition, whereas network segregation, or lower between-network connectivity and higher within-network connectivity, is important for sustained attention. As mentioned above, the go/no-go task is thought to index both response inhibition and sustained attention (Simmonds et al. 2008; Smith et al. 2013), yet it remains unknown how brain network organization supports go/no-go task performance in children, particularly because cognitive strategies might differ throughout development and into adulthood (Munakata et al. 2012; Chevalier et al. 2015; Niebaum and Munakata 2023). As response inhibition and sustained attention are associated with different patterns of network integration and segregation, by quantifying these different aspects of topology during the performance of a go/no-go task in children, we can begin to disentangle how these two cognitive strategies are employed in youth.

### The influence of rewards on response inhibition and sustained attention

Additionally, performance during tasks probing response inhibition (Geier and Luna 2009; Burton et al. 2021) and sustained attention (Smith et al. 2011; Fortenbaugh et al. 2017) often improves when rewards are offered for good performance. This is the case for performance on go/no-go tasks in children and adolescents (Winter and Sheridan 2014; Demurie et al. 2016; Miyasaka and Nomura 2019). Rewards have been shown to elicit stronger, more proactive, and more consistent engagement of task-relevant brain regions and networks, including the fronto-parietal and visual networks (Smith et al. 2011; Etzel et al. 2016; Esterman et al. 2017; Shashidhara et al. 2019). Rewards also influence functional connectivity. For example, in adults, when monetary rewards are provided for performance during a cognitive control task, functional connectivity strength increases between pairs of regions that span cognitive control-related (e.g., fronto-parietal and cingulo-opercular) and reward networks (Dixon and Christoff 2012; Boehler et al. 2014; Teslovich et al. 2014; Bahlmann et al. 2015; Cubillo et al. 2019). Additionally, in adolescents, reward-driven increases in connectivity within the salience network are associated with improvements in accuracy during a rewarded antisaccade task (Hallquist et al. 2018). However, it remains unknown how rewards shape large-scale functional network organization in the service of response inhibition and sustained attention in children.

### The current study

Here, we investigated differences in functional brain network organization between the resting state and go/no-go tasks with and without monetary rewards for performance, in a sample of 26 children between the ages of 8 and 12 years. With this age range, we are able to probe brain network organization when response inhibition and sustained attention are established but not yet fully developed (Lin et al. 1999; Betts et al. 2006; Best and Miller 2010; Luna et al. 2013; Brydges et al. 2014). We characterized brain network organization across the whole brain and as related to key networks, including those implicated in cognitive control (fronto-parietal, cingulo-opercular), other aspects of higher-order functioning (salience, default mode, dorsal and ventral attention), rewards, and sensorimotor processing (motor, visual). We further examined specific regions found to be relevant to response inhibition, attention, and reward processing in prior literature and assessed the roles of these regions within the larger brain network. Our primary goal was to determine whether functional brain networks reconfigure in a way that is more consistent with response inhibition (increased integration) or sustained attention (decreased integration) during the go/no-go task in children, thus identifying the predominant neurocognitive strategy for go/no-go task execution within this age range. The inclusion of the rewarded go/no-go task allowed us to test how the pattern of brain network reconfiguration shifts in the presence of rewards for good task performance. We expected that with the addition of rewards brain network organization would become even more optimal for a given strategy. The results from this study improve the general understanding of how functional brain network organization underlies response inhibition, sustained attention, and reward processes in late childhood.

## Materials and Methods

### Participants

Participants were 30 typically-developing children between the ages of 8 and 12 years. Participants were enrolled as part of a larger study and, as relevant here, completed functional magnetic resonance imaging (fMRI) scans and self-report questionnaires. Four participants were excluded from the study for the following reasons: task administration error (n = 2), failure to pass cortical segmentation quality check due to image intensity abnormalities (n = 1), and incidental findings during the MRI scan (n = 1). Therefore, a total of 26 participants (11 females, *mean* = 10.42 years, *sd* = 1.47 years) were included in analyses. See **Table 1** for demographic information, including age, sex, race, family income, and parental education.

**Table 1.** Demographic Characteristics.

| Variable | Mean±SD |
| --- | --- |
| Age (years) | 10.42±1.47 |
| Sex (M/F) | N=11 (42.3%) / N=15 (55.7%) |
| Estimated IQ<sup>a</sup> | 116.88±12.26 |
| Word Reading<sup>b</sup> | 115.84±10.57 |
| Pubertal development stage<sup>c</sup> |  |
| Prepubertal | 15 (57.7%) |
| Early puberty | 7 (26.9%) |
| Middle puberty | 3 (11.5%) |
| Late puberty | 1 (3.8%) |
| Race |  |
| Asian | 2 (7.7%) |
| Black/African American | 1 (3.8%) |
| White/Caucasian | 21 (80.8%) |
| Multiracial | 2 (7.7%) |
| Family Income<sup>d</sup> |  |
| Up to \$24,999 | 1 (3.8%) |
| \$25,000 - \$74,999 | 2 (7.7%) |
| Above \$75,000 | 20 (77.0%) |
| Parental Education<sup>d</sup> |  |
| High school diploma/GED | 1 (3.8%) |
| Bachelor's degree | 6 (23.1%) |
| Master's degree | 8 (30.8%) |
| Doctoral or professional | 11 (42.3%) |

Total sample: N = 26
<sup>a</sup> Estimated intelligence quotient (IQ) determined using the Wechsler Intelligence for Children, Fifth Edition (WISC-V); one participant did not complete the WISC-V (M, 10.4 years).
<sup>b</sup> Word reading (standard score) determined using the Wechsler Individual Achievement Test, 3rd Edition (WIAT-III); one participant did not complete the WIAT-III (M, 10.4 years).
<sup>c</sup> Pubertal development stage determined using parent-report on the Pubertal Development Scale (Petersen et al. 1988).
<sup>d</sup> Family income and parental education established using the MacArthur Scale (Adler et al. 2000) administered to parents; three parents declined to answer the family income question; all parents answered the parental education question.

Study procedures were reviewed and approved by the University of North Carolina at Chapel Hill Institutional Review Board. Participants were recruited through flyers, website or magazine advertisements, and listserv emails. Potential participants were excluded if they had an intelligence quotient (IQ) less than 80 as measured by the *Wechsler Intelligence Scale for Children, Fifth Edition* (WISC-V; Wechsler, 2014), a standard score less than 85 on the Word Reading subtest of the *Wechsler Individual Achievement Test, Third Edition* (WIAT-III; Wechsler, 2002), a vision or hearing impairment, an MRI contraindication (e.g., implanted electrical devices, dental braces), or were diagnosed with a psychiatric, neurologic or learning disorder. The *Diagnostic Interview Schedule for Children - Version IV*, *Parent Report* (DISC; Shaffer et al. 2000) was used to assess the presence of psychiatric disorders. Pubertal stage was assessed using the parent-report version of the Pubertal Development Scale (Petersen et al. 1988).

### Procedures

The current study was conducted over the course of three separate visits. Participants first completed a behavioral visit, followed by two MRI scanning sessions. At the behavioral visit, parents provided written consent and participants provided written assent. Additionally, the behavioral visit included cognitive and achievement testing, self-report questionnaires, and an MRI mock scan. During the mock scan, participants lay inside a replica of an MRI scanner while wearing a motion-tracking headband. Participants were exposed to scanner noises, practiced the standard and rewarded go/no-go tasks, and received real-time feedback on their movement. The mock scan session allowed participants to practice being still while becoming familiar with the scanner environment.

Participants completed two identical scanning sessions an average of 12.3 days apart (range = 3-42 days). Both MRI scanning sessions included an MPRAGE anatomical T1-weighted scan, two resting state scans (5 minutes each), two standard go/no-go task scans (6.5 minutes each), and four rewarded go/no-go task scans (6 minutes each). Participants were provided with a weighted blanket, headphones, and padding for comfort. A handheld button box was used to record participant responses. Visual task stimuli were projected onto a screen mounted to the inside of the magnet bore. During the resting state scans, participants were instructed to remain awake with their eyes open and fixate on a white central cross on a gray background. Tasks were presented using PsychoPy v1.85.1 (Peirce 2007; Peirce 2009).

The measures collected in this study were a subset of measures collected from the larger study; only measures relevant to this study will be described below.

### Go/no-go tasks

Two versions of a go/no-go task were administered to participants during the MRI scan: a standard go/no-go task and a rewarded go/no-go task.

During the standard go/no-go task, eight sports balls were presented on the screen one at a time (**Figure 1a**). Two of the eight sports balls were randomly selected, separately for each participant, as ‘no-go’ stimuli. The remaining 6 sports balls were ‘go’ stimuli. Participants were instructed to press a button with the index finger of their right hand to indicate when ‘go’ sports balls were presented (∼73% of trials) and to withhold from pressing a button when ‘no-go’ sports balls were presented (∼27% of trials). Participants were instructed to respond to ‘go’ sports balls as quickly as possible. There were 128 trials in each of two standard go/no-go runs. Stimulus order was pseudorandom such that there were between two and four consecutive go trials before each no-go trial, with 16 instances of two consecutive go trials, 10 instances of three consecutive go trials, and eight instances of four consecutive go trials during each run. Stimuli were displayed for 600ms with a jittered interstimulus interval ranging from 1.25 to 3.25s selected from a uniform distribution. Participants could respond to the stimulus through the end of the interstimulus interval.

**Figure 1.**
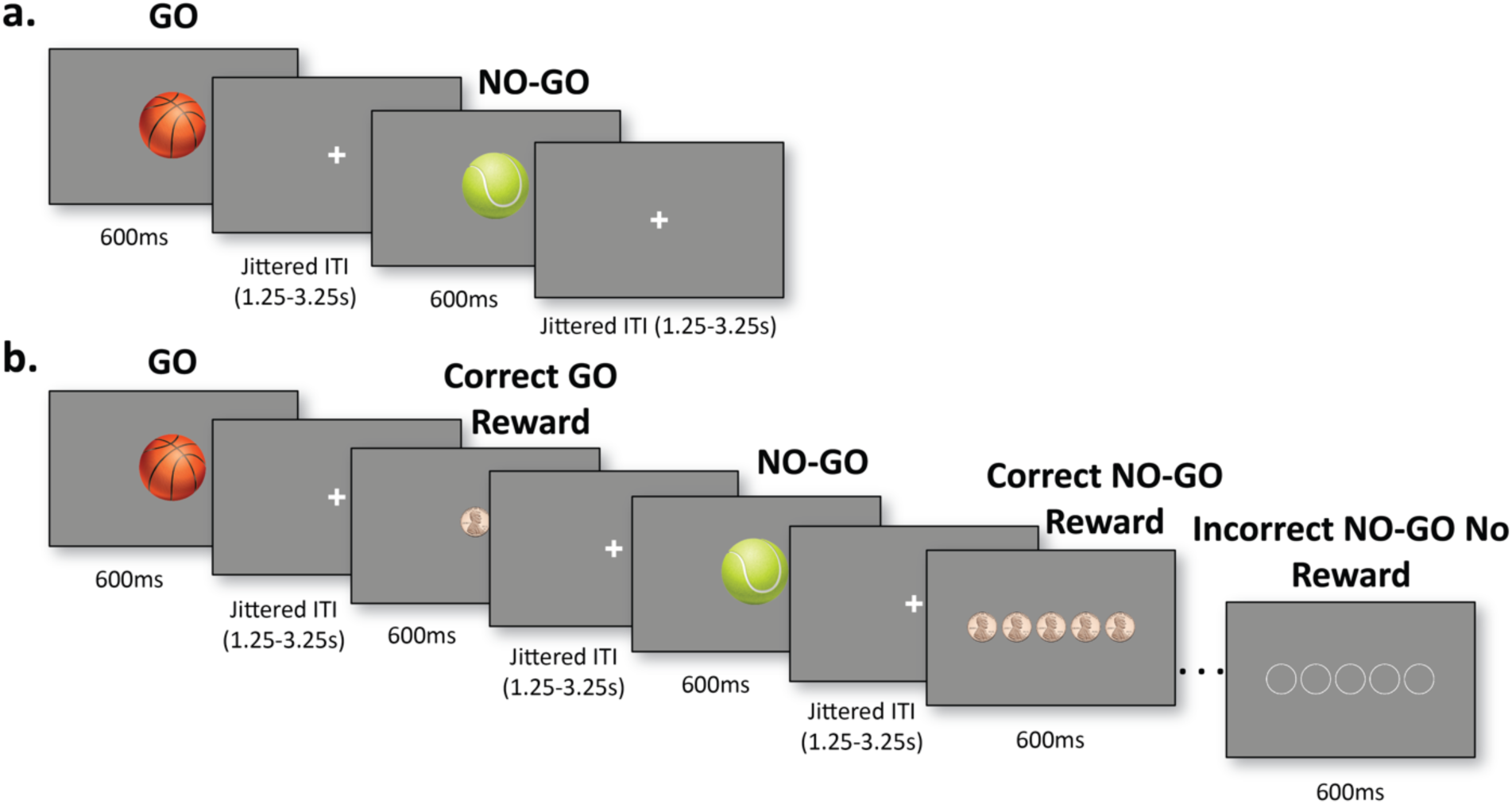
fMRI Task stimuli. **a.** Standard go/no-go task: Sports balls were presented one at a time. In this example, basketballs were ‘go’ stimuli and the correct response was a button press. Tennis balls were ‘no-go’ stimuli, and the participant was expected to withhold pressing the button. **b.** Rewarded go/no-go task: Each sports ball stimulus was followed by the reward that was earned (pennies shown) or not earned (empty circles shown). The potential reward for each go trial was one penny. The potential reward for each no-go trial was five pennies.

During the rewarded go/no-go task, the same sports ball stimuli were used and participants were given the same instructions as in the standard go/no-go task (**Figure 1b**). They were additionally told that correct, fast responses on go trials (≤650 ms) and correct non-responses on no-go trials would be rewarded. There were 64 trials in each of four rewarded go/no-go runs. Stimulus order and timing were identical to those of the standard go/no-go task, with the following modifications: 1) the interval between the stimulus and the reward feedback was jittered identically to the interstimulus interval between trials; and 2) reward feedback following the jitter was presented for 600ms seconds. Participants were visually presented with a reward of one penny for fast, correct responses on go trials and five pennies for correctly withheld responses on no-go trials. Participants viewed the same number of empty circles on trials that were not rewarded. Participants accumulated money throughout the task runs and were given the earned amount at the end of the visit.

Behavioral performance on the standard and rewarded go/no-go tasks was indexed by commission error rate and tau. Commission error rate was calculated as the proportion of no-go trials on which the participant failed to inhibit the button press and was used as an index of response inhibition. Tau quantifies the positive skew of the response time distribution when fitted with an exponential-Gaussian (ex-Gaussian) curve (Balota and Yap 2011) and is used to represent the frequency of longer response times. These long response times are thought to be due to lapses in attention (Leth-Steensen et al. 2000). Thus, tau was used as an index of sustained attention. The timefit function from the *retimes* package in R was used to bootstrap the correct go trial response times with 5,000 iterations of sampling with replacement to achieve a sample of the same size as the original response time vector in each iteration. Parameters were estimated in each iteration and then tau, which reflects the mean and standard deviation of the exponential component, was quantified through maximum likelihood estimation (Massidda 2013). Commission error rate and tau were normally distributed on both tasks (all Shapiro-Wilk test of normality *p*-values > .096).

Individual task runs were excluded from analyses if there was greater than 20% omission rate on go trials. Across all participants and both fMRI sessions, six runs were excluded from the standard go/no-go task (two runs from each of two participants and one run from each of two participants) and seven runs were excluded from the rewarded go/no-go task (three runs from one participant and one run from each of four participants). Additionally, behavioral data was not logged for one standard go/no-go run for one participant and one rewarded go/no-go run for one participant, thus those runs were excluded as well. These participants were still included in analyses as they had at least one good run of data.

Additionally, rewarded go/no-go data was not collected from one participant (male, 10.4 years), thus this participant was not included in analyses involving the rewarded go/no-go task. This resulted in *N* = 26 participants included in analyses with the standard go/no-go task data and *N* = 25 participants included in analyses with the rewarded go/no-go task data based on behavioral criteria.

### Reward responsivity

Participants completed the Behavioral Inhibition System and Behavioral Activation System scales (BIS/BAS; Carver and White 1994). The BIS/BAS measures inhibitory and approach motivation systems, including reward responsiveness, which is assessed with five items probing response to rewarding events. The mean scale BAS reward responsiveness score was used in analyses. Internal consistency for the BAS reward responsiveness scale was *α* = .72.

### fMRI data acquisition

Neuroimaging data were collected with a 32-channel head coil on a 3-tesla Siemens MAGNETOM Prisma-fit whole-body MRI machine at the University of North Carolina at Chapel Hill Biomedical Research Imaging Center. A high-resolution anatomical T1-weighted MPRAGE scan was collected (TR = 2400 ms, TE = 2.22 ms, FA = 8°, field of view 256mm x 256mm, 208 slices, in-plane voxel size: 0.8 mm). Whole brain functional data were collected with a gradient-echo T2*-weighted EPI sequence (39 slices, TR: 2000 ms, TE: 25 ms, flip angle: 77°, echo spacing: 0.54 ms, field of view: 230 × 230 mm, voxel dimensions: 2.9 mm × 2.9 mm × 3 mm). A total of 300 timepoints were collected during the resting state (150 timepoints and 5 minutes per run), 390 timepoints during the standard go/no-go task (195 timepoints and 6.5 minutes per run), and 740 timepoints during the rewarded go/no-go task (185 timepoints and 6.17 minutes per run).

### fMRI data processing

fMRI data preprocessing was performed using fMRIPrep 1.5.0 (Esteban et al. 2018), which includes EPI to T1w coregistration, susceptibility artifact correction, normalization to MNI space, and estimation of motion parameters. Details can be found in the **Supplementary Materials**. Following minimal preprocessing by fMRIPrep, several functional connectivity (FC) postprocessing steps were applied following FC processing guidelines <u>(Ciric et al. 2017</u>; see **Supplementary Materials**). We applied 32 parameter nuisance regression (6 degrees of motion, white matter and CSF signal, with temporal, quadratic, and quadratic temporal derivatives) along with bandpass spectral filtering (0.009 - 0.08 mHz). Global signal was not included as a regressor due to concerns that global signal regression alters the functional network structure of the brain graphs (Murphy et al. 2009; Saad et al. 2012; Caballero-Gaudes and Reynolds 2017; Li et al. 2019). Bandpass filtering was also applied to the regressor matrix (Hallquist et al. 2013; Lindquist et al. 2019). As scanner-estimated motion can be inflated as a result of respiration-related magnetic field disruptions, notch-filtered framewise displacement (FD; <u>Power et al. 2012)</u> was used for scrubbing (Fair et al. 2020; Gratton et al. 2020) with a frequency band of 0.31-0.41 mHz. Scrubbing was performed with a 0.2mm notch-filtered FD threshold, with five contiguous timepoints required to be included in the timeseries. To combine scrubbing and spectral filtering, spectral interpolation based on the XCP pipeline (Ciric et al. 2018) was used. All postprocessing steps were implemented using a customized processing pipeline (clpipe; Henry et al. 2023). Additionally, in an effort to further reduce noise contamination, rest or task runs that had a mean notch-filtered FD exceeding 0.5mm were excluded.

Runs with poor fMRI data quality (e.g., insufficient coverage, poor registration) were excluded from analyses. Across all participants and both fMRI sessions, one run was excluded from the standard go/no-go task and six runs were excluded from the rewarded go/no-go task (three runs from one participant and one run from each of three participants) for poor fMRI data quality.

As reliability of functional connectivity estimates increase with the amount of within-participant data (Nee 2019; Noble et al. 2019), we combined data across both sessions within each participant and each task to maximize the timepoints for each go/no-go fMRI task. At least 50 good timepoints for each run, as well as at least 150 good timepoints across runs for each cognitive state, were required to be included in analyses. This resulted in one run of the rewarded go/no-go task excluded from one participant. As only two runs of the rewarded go/no-go task were collected from this participant (female, 9.6 years) and the other run was excluded for poor quality (see prior paragraph), this participant was excluded from analyses involving the rewarded go/no-go task. Additionally, as mentioned above (see *Go/no-go tasks* section), data from the rewarded go/no-go task was not collected from one participant. Therefore, the final sample included *N* = 26 participants in analyses involving the resting state and standard go/no-go task, but *N* = 24 participants in analyses involving the rewarded go/no-go task. For more information on the number of runs per participant and per cognitive state see **Table 2**.

**Table 2.** Runs and timepoints retained in analyses by cognitive state.

|  | Number of runs retained | Number of timepoints retained | Minutes retained |
| --- | --- | --- | --- |
| **Resting state** | 4 runs (N = 24)<br>2 runs (N = 2) | 219 – 599 timepoints<br>mean = 537 (sd = 96.42) | 7.3 – 20.0 minutes<br>mean = 17.90 (sd = 3.21) |
| **Standard go/no-go** | 4 runs (N = 18)<br>3 runs (N = 3)<br>2 runs (N = 5) | 247 – 756 timepoints<br>mean = 602.88 (sd = 156.90) | 8.23 – 25.20 minutes<br>mean = 20.10 (sd = 5.23) |
| **Rewarded go/no-go** | 8 runs (N = 10)<br>7 runs (N = 9)<br>6 runs (N = 1)<br>5 runs (N = 2)<br>4 runs (N = 1)<br>3 runs (N = 1)<br>0 runs (N = 2) | 478 – 1442 timepoints<br>mean = 1108.33 (sd = 260.42) | 15.93 – 48.07 minutes<br>mean = 36.94 (sd = 8.68) |

For the resting state and standard go/no-go task all participants were included (n = 26). For the rewarded go/no-go task, sufficient data was not collected from two participants (n = 24). See *Go/no-go Tasks* and *fMRI Data Processing* for descriptions of subjects and runs removed from the analyses.

### Functional connectivity

BOLD timeseries were extracted from 300 spherical 8mm-diameter regions of interest (ROIs; Seitzman et al. 2020; **Figure 2a**) using clpipe (Henry et al. 2023). This atlas includes 13 functional networks, as well as a group of regions unassigned to a network. Due to a limited field of view, data was not collected in all participants and all fMRI scans from two ventral regions of the cerebellum, one region assigned to the cingulo-opercular network and the other assigned to the dorsal attention network. Thus, a final set of 298 ROIs was used.

**Figure 2.**
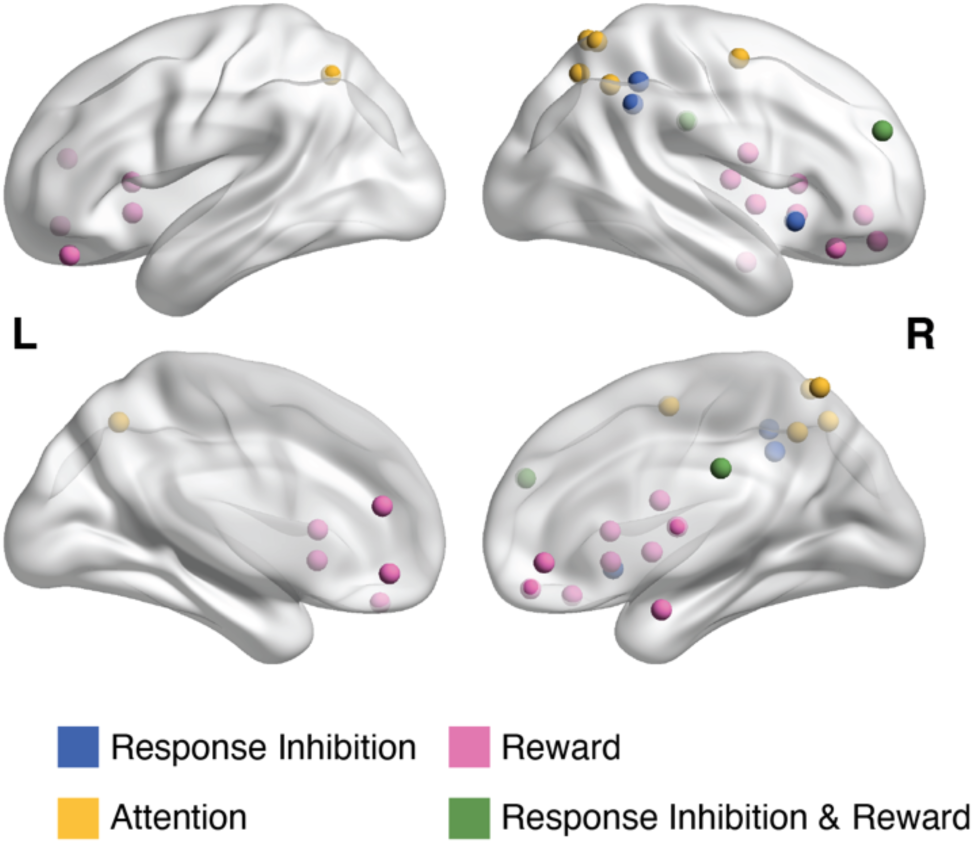
Neurosynth-derived regions of interest corresponding to the search terms “response inhibition” (3), “attention” (6), “reward” (14), and both “response inhibition” and “reward” (2). Image generated in BrainNet Viewer (Xia et al. 2013).

We conducted both whole brain and network-level analyses. For network analyses, we were specifically interested in nine networks implicated in task-relevant cognitive functioning (i.e., fronto-parietal [36 ROIs], cingulo-opercular [29 ROIs], salience [13 ROIs], default mode [65 ROIs], dorsal attention [15 ROIs], ventral attention [11 ROIs], reward [8 ROIs], dorsal somatomotor [40 ROIs], and visual [37 ROIs] networks).

Subsequent steps were conducted in R (R Core Team 2019; RStudio Team 2019). To maximize the number of timepoints entered into each functional connection, the timeseries were concatenated across runs and sessions within each cognitive state (see **Table 2**). Each run was detrended and standardized to control for differences in BOLD intensity across runs prior to concatenation. FC between each pair of ROIs was estimated using Pearson’s correlation coefficient. This resulted in a 298 x 298 correlation matrix for each participant and for each cognitive state (i.e., resting state, standard go/no-go, rewarded go/no-go). Correlation matrices were Fisher z-transformed prior to the construction of brain graphs.

### Graph construction

The Fisher z-transformed correlation matrices were thresholded using an absolute correlation threshold to remove negative correlations and low magnitude correlations that may be spurious. Given that there is no universal benchmark for threshold specification, we ran analyses across a range of FC thresholds, from *r* = 0.20-0.45 in increments of 0.05. These FC thresholds approximate common brain graph densities reported consistently in the literature (2-25%) (Power et al. 2011; Cohen and D’Esposito 2016; Gordon et al. 2018; Le et al. 2020; Seitzman et al. 2020; Demeter et al. 2023). We averaged the proportion of disconnected nodes in each network of interest in each cognitive state at each threshold. An elbow point was estimated from the distribution of disconnected nodes across each threshold using the *akmedoids* package in R (Adepeju et al. 2020), in order to select the greatest threshold before a sharp increase in disconnected nodes. The elbow point was estimated to fall between the *r* = 0.30 and *r* = 0.35 thresholds (**Supplementary Figure 2**). Thus, the FC thresholds included in the final analyses were 0.20, 0.25, and 0.30. All remaining edges after thresholding retained their weight.

### Graph metrics

Weighted metrics quantifying network integration and network segregation were calculated from the brain graph for each participant and each cognitive state using the *igraph* and *braingraph* packages in R (Csardi and Nepusz 2006; Watson 2020). We focused on the whole-brain metrics of global efficiency and modularity to quantify whole-brain integration and whole-brain segregation, respectively. Global efficiency is a measure of information transfer across the whole brain system without regard for network membership, and thus is a measure of whole-network integration. Global efficiency was calculated as the inverse of the average shortest path length (Latora and Marchiori 2001; Watson 2020). Modularity is a measure of the degree to which the brain segregates into distinct subnetworks, or communities, with many/stronger connections within these subnetworks and fewer/weaker connections between subnetworks (Bullmore and Bassett 2011). Modularity was calculated using the preassigned network membership from the functional atlas (Csardi and Nepusz 2006; Seitzman et al. 2020). Greater values of modularity indicate that the whole brain system is more segregated.

To characterize between-network integration, we calculated node dissociation index (Cary et al. 2016). Node dissociation index was calculated for each node as the sum of weighted connections to nodes in every network other than its own divided by the total weighted connections. When averaged across nodes in each network, node dissociation index quantifies how much a given network is integrated with other networks. To characterize within-network coherence, we calculated an unstandardized, weighted variant of within-module degree (Guimerà and Nunes Amaral 2005). Within-module degree was calculated for each node as the sum of weighted connections with all other nodes in the same network. When averaged across all nodes within a network, within-module degree quantifies how strongly interconnected the nodes within a given network are with each other. Unstandardized within-module degree was calculated so that an average across all nodes within a network could be calculated. Finally, density and mean FC of the whole brain were calculated. Density was calculated as the ratio of edges included in the graph after thresholding to the number of possible edges (Wasserman and Faust 1994). Mean FC was calculated as the average of all edge values in the matrix remaining after thresholding. Brain metrics were calculated at each threshold (*r*-values = 0.20, 0.25, 0.30) and averaged across thresholds prior to analysis as in other studies (Cohen and D’Esposito 2016; Gordon et al. 2018; Demeter et al. 2023).

### Meta-analytic ROI identification and nodal analyses

ROIs were identified through meta-analytic association test maps downloaded from Neurosynth (https://neurosynth.org/) (Yarkoni et al. 2011) in early November 2023 for the following terms: “response inhibition”, “attention”, and “reward”. The term “attention” was used instead of “sustained attention” as the map for the term “sustained attention” was largely empty. We suspect that the relative vagueness in the definition and use of the term “sustained attention” (Oken et al. 2006) may have contributed to the lack of unique, consistent activations in the Neurosynth association test map (Shine and Poldrack 2018). The 300 ROIs from the Seitzman atlas (Seitzman et al. 2020) were overlayed on each association map and ROIs with at least 50% overlap were identified as specific to response inhibition, attention, and/or reward.

Twenty five regions were identified from the association maps: three response inhibition regions, six attention regions, fourteen reward regions, and two regions that were present in both the response inhibition and reward maps (see **Table 3** and **Figure 2**).

**Table 3.**
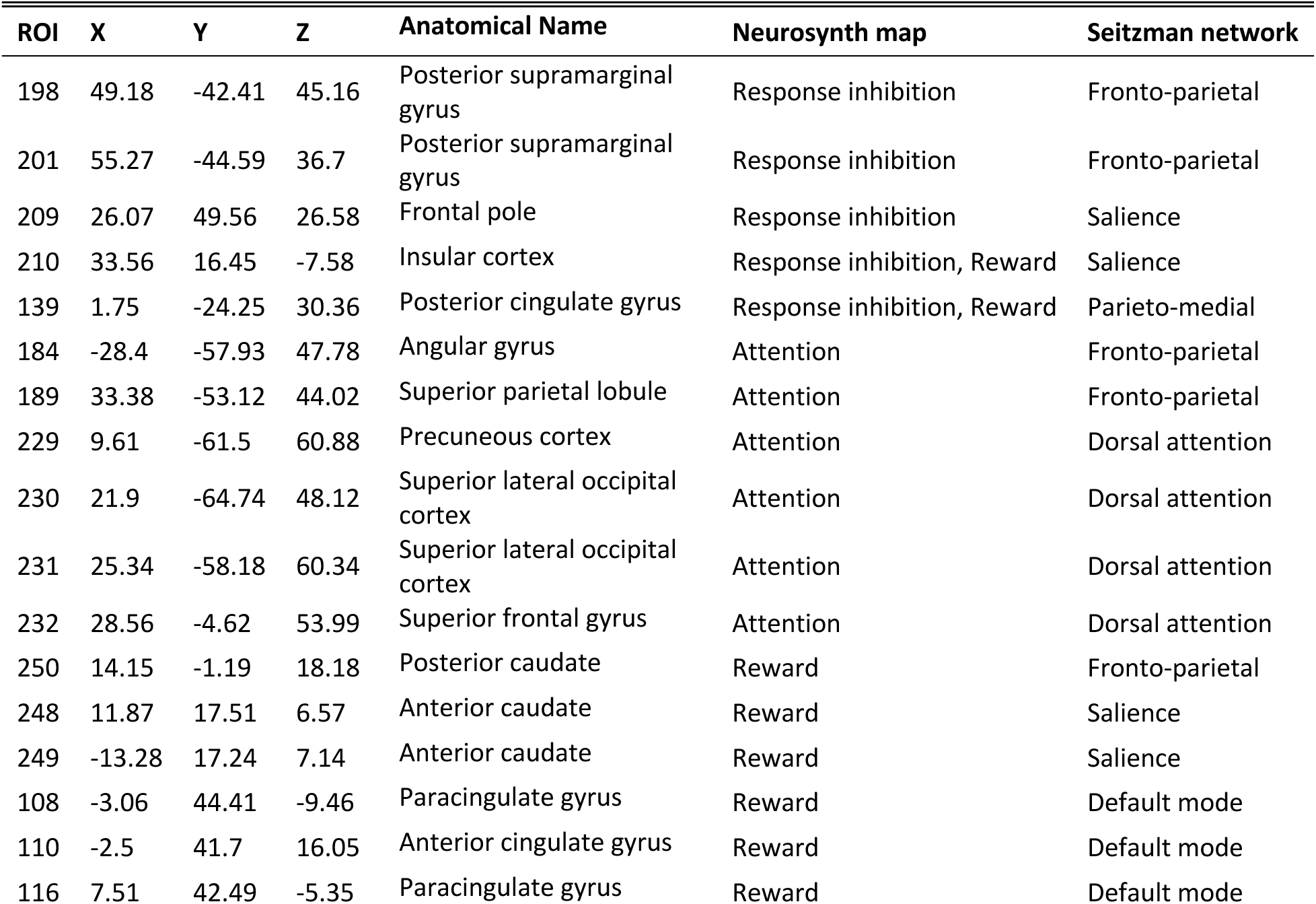
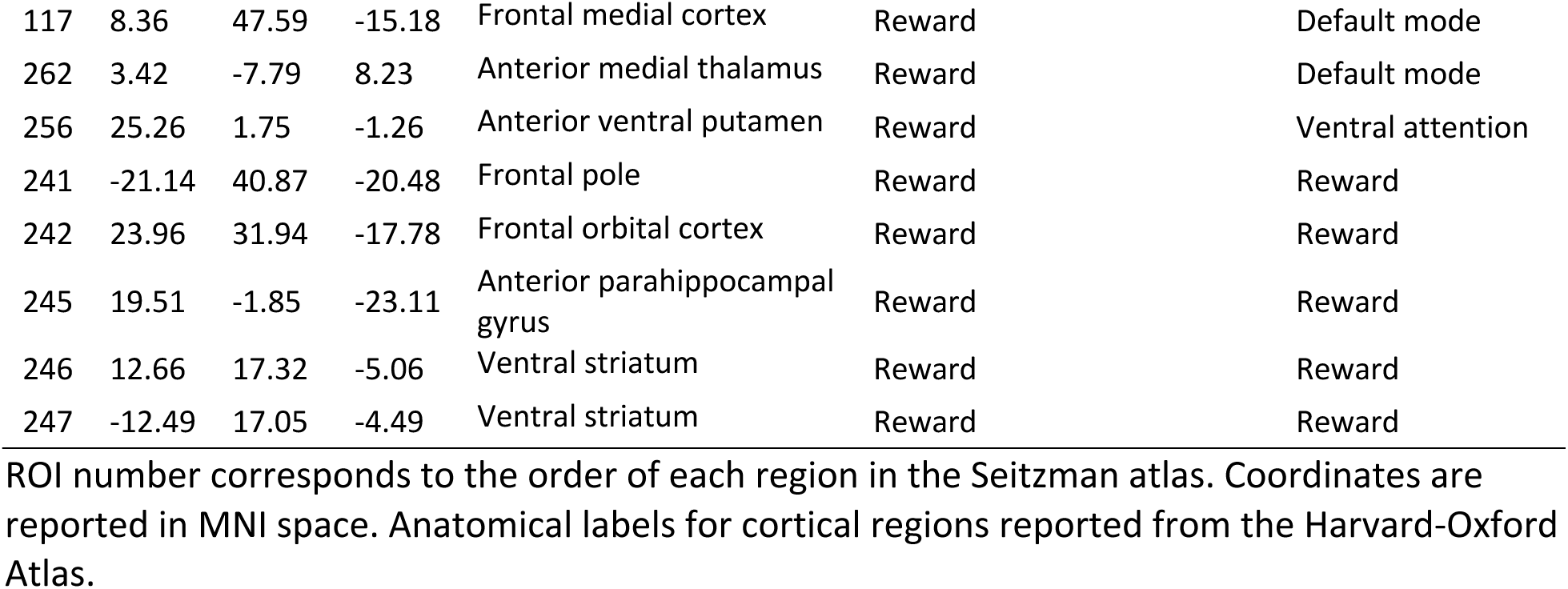
Seitzman atlas regions overlapping with Neurosynth association maps.

### Analyses

Paired t-tests were conducted to investigate differences in behavioral task performance (i.e., commission error rate and tau) between the standard and rewarded go/no-go tasks. Results were corrected for two tests with the Benjamini-Hochberg false discovery rate (FDR) at *q* < .05 (Benjamini and Hochberg 1995). A post hoc analysis was run to assess if changes in task performance between the standard and rewarded go/no-go task were related to reward responsivity, measured on the BIS/BAS (Carver and White 1994).

Next, we quantified how brain organization was different across pairs of cognitive states: rest versus standard go/no-go and standard go/no-go versus rewarded go/no-go. A mixed effects model was conducted to compare graph metrics across each pair of cognitive states using contrast coding and the following equation:

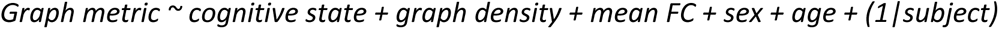

The contrast to test for differences between the resting state and the standard go/no-go task was [-1 1 0] for rest, standard go/no-go, and rewarded go/no-go respectively. The contrast to test for differences between the standard and rewarded to/no-go task states was [0 -1 1] for rest, standard go/no-go, and rewarded go/no-go respectively. Graph density, mean FC, sex, and age were included in the models as fixed effects and a random participant-level intercept was used to control for inter-individual variation. Mean FC and graph density were controlled for given that differences in mean FC and graph density impact estimation of network statistics (van Wijk et al. 2010; van den Heuvel et al. 2017; Hallquist and Hillary 2019). We first related whole brain organization (global efficiency, modularity) across each pair of states. We FDR-corrected at *q* < .05 for the two graph metrics separately for each cognitive state pair. Then, we related network-level metrics (node dissociation index and within-module degree for each of the nine networks of interest) across each pair of states. For network-level comparisons, we FDR-corrected at *q* < .05 for the 18 network-level metrics separately for each cognitive state pair.

Finally, we quantified how nodal properties (node dissociation index and within module degree) differed across the same pairs of cognitive states for the 25 Neurosynth-derived ROIs. Nodal properties were calculated using the Seitzman atlas network assignments. The same mixed effects models, contrast coding, and covariates were used as in the above analysis. We FDR-corrected at q < .05 for the 50 nodal-level metrics separately for each cognitive state pair.

For all analyses, we report standardized betas (*β*) to allow for more direct interpretation of effect sizes.

## Results

### Behavioral results

First, we assessed task performance during the standard and rewarded go/no-go tasks (**Table 4**). Between the standard and rewarded go/no-go tasks there was a significant decrease in commission error rate (*t*(23) = 2.46, adjusted-*p* = .043), but no change in tau (*t*(23) = 1.64, adjusted-*p* = .115). In the standard go/no-go task, commission error rate and tau were not significantly related (*r* = .27, *p* = .179). However, in the rewarded go/no-go task, commission error rate and tau were significantly related (*r* = .46, *p* = .024; **Table 4**).

**Table 4.**
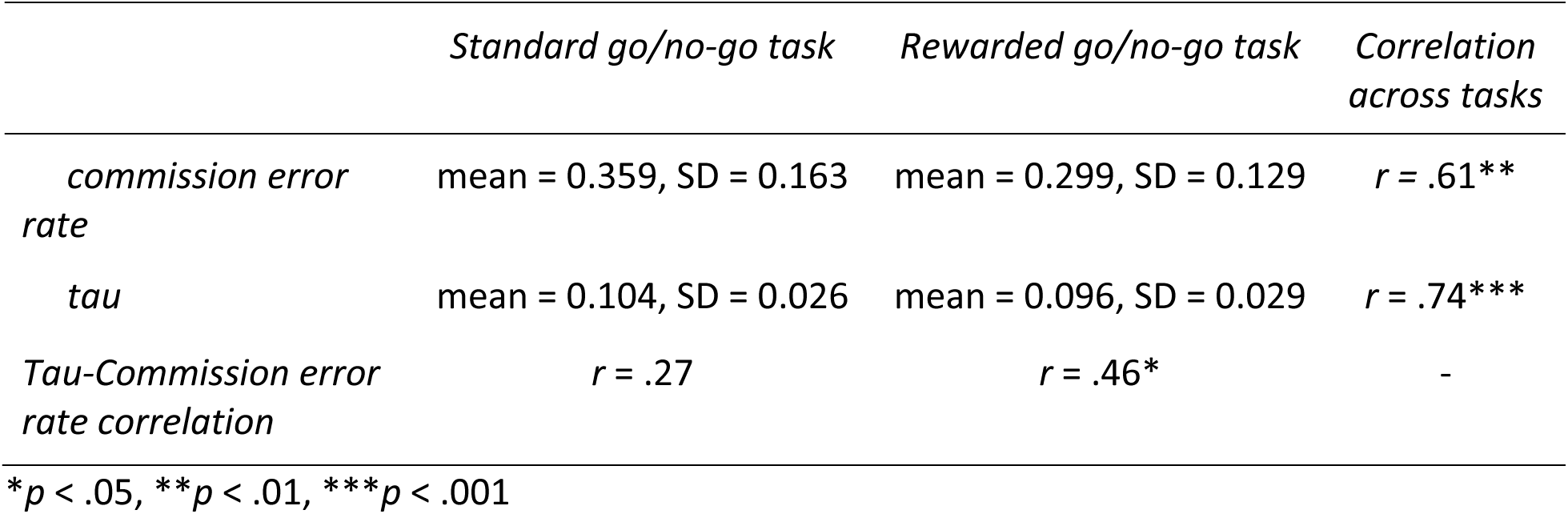
Behavioral measures of task performance.

|  | *Standard go/no-go task* | *Rewarded go/no-go task* | *Correlation across tasks* |
| --- | --- | --- | --- |
| *commission error rate* | mean = 0.359, SD = 0.163 | mean = 0.299, SD = 0.129 | $r = .61^{**}$ |
| *tau* | mean = 0.104, SD = 0.026 | mean = 0.096, SD = 0.029 | $r = .74^{***}$ |
| *Tau-Commission error rate correlation* | $r = .27$ | $r = .46^*$ | - |

\* $p$ < .05, \*\* $p$ < .01, \*\*\* $p$ < .001

As past literature has shown improvements in sustained attention with the addition of rewards (Smith et al. 2011; Fortenbaugh et al. 2017), it is surprising that we did not see improvements in our behavioral measure of sustained attention in the rewarded go/no-go task. It is possible that individual differences in sensitivity to rewards could impact the degree of improvement across participants. To test this possibility, we ran a post hoc analysis to investigate if self-reported reward responsivity on the BIS/BAS (mean = 3.41, SD = 0.45) related to change in each test performance measure separately. In linear regressions controlling for age and sex, we found that reward responsivity was significantly related to change in commission error rate (*β* (standardized beta) = -0.43, *p* = .042), and marginally related to change in tau (*β* = -0.42, *p* = .054).

**Figure 2.**
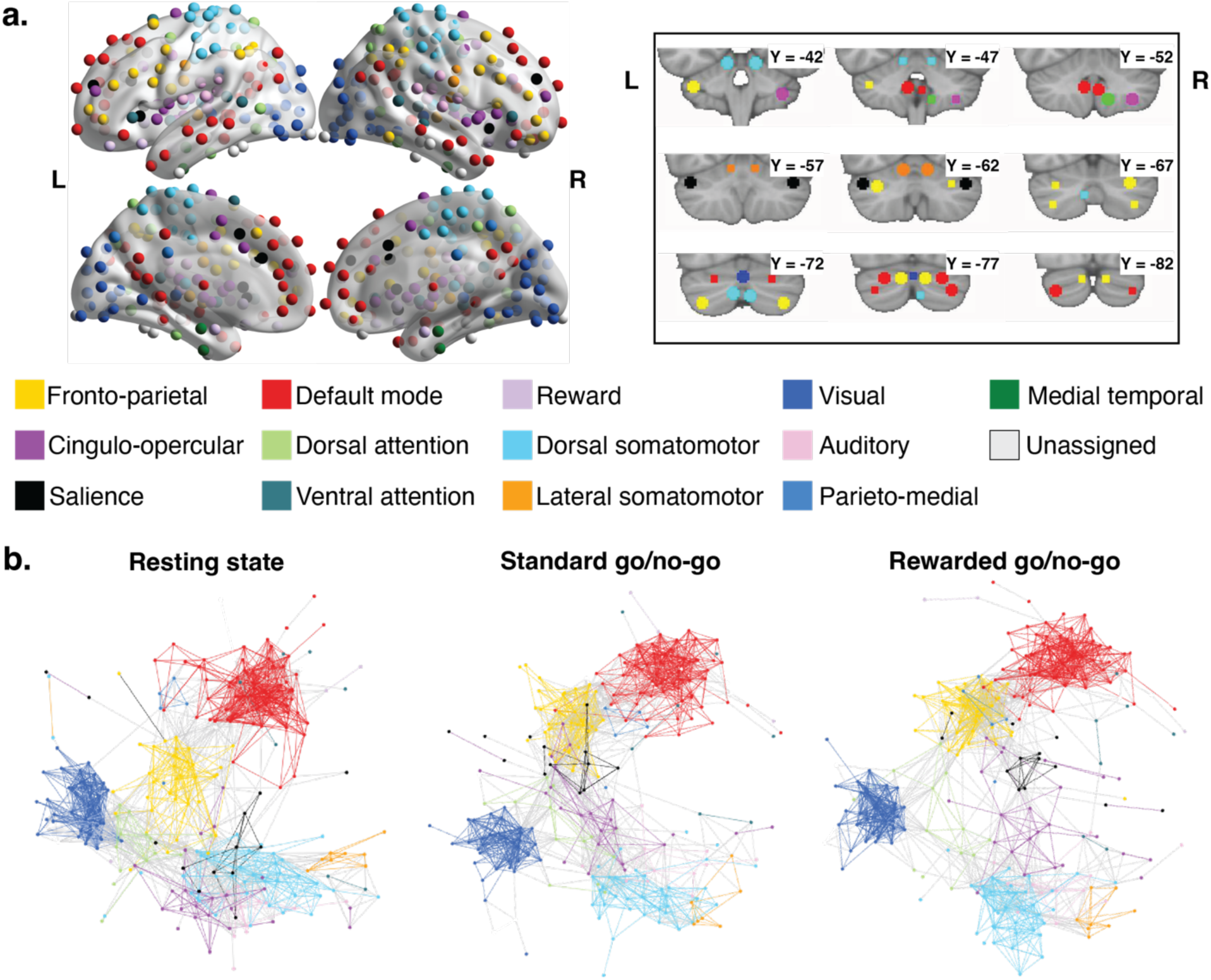
Functional brain atlas and brain graphs for each cognitive state. **a.** 298 regions of interest (ROIs) (Seitzman et al. 2020). Cortical and subcortical regions (n = 273) are depicted on medial and lateral views of a 3D brain on the left (images generated in BrainNet Viewer; Xia et al. 2013). Serial coronal views of the cerebellar regions (n = 25) are in the box to the right (images generated in FSLview; Jenkinson et al. 2012). Figure adapted from Seitzman et al. (2020). **b.** Spring graphs for each cognitive state were calculated from the average cognitive state correlation matrix across all participants. ROIs are depicted as nodes. Correlations surviving thresholding at *r*=0.25 are depicted as edges. Within-network edges are colored to correspond with the network color, while between-network edges are colored gray. Spring graphs were generated in Cytoscape (Shannon et al. 2003) using the edge-weighted spring embedded layout.

### Differences in brain network organization between cognitive states

First, we characterized brain network organization during the resting state, the standard go/no-go task, and the rewarded go/no-go task (**Figure 2b**). To quantify differences in whole brain organization between pairs of cognitive states, we estimated global efficiency and modularity. During the standard go/no-go task as compared to the resting state, we observed significantly decreased global efficiency (*β* = -0.26, adjusted-*p* < .001) and increased modularity (*β* = 0.17, adjusted-*p* = .008). During the rewarded go/no-go task as compared to the standard go/no-go task, we observed significantly decreased global efficiency (*β* = -0.32, adjusted-*p* < .001) and increased modularity (*β* = 0.36, adjusted-*p* < .001; **Supplementary Table 1**).

To quantify differences in network-level organization between pairs of cognitive states, we estimated between-network integration (node dissociation index) and within-network coherence (within-module degree) for each network of interest. During the standard go/no-go task as compared to the resting state, we observed significantly decreased node dissociation index in the fronto-parietal (*β* = -0.29, adjusted-*p* < .001), cingulo-opercular (*β* = -0.26, adjusted-*p* = .036), salience (*β* = -0.25, adjusted-*p* = .032), default mode (*β* = -0.23, adjusted-*p* < .003), reward (*β* = -0.42, adjusted-*p* = .011), and dorsal somatomotor (*β* = -0.42, adjusted-*p* < .001) networks. During the standard go/no-go task, we also observed significantly increased within-module degree in the fronto-parietal network (*β* = 0.38, adjusted-*p* < .001) and significantly decreased within-module degree in the visual network (*β* = -0.28, adjusted-*p* = .005). All other adjusted-*p*s > .061 (**Supplementary Table 2**).

During the rewarded go/no-go task as compared to the standard go/no-go task, we observed significantly decreased node dissociation index in the fronto-parietal (*β* = -0.39, adjusted-*p* < .001), cingulo-opercular (*β* = -0.34, adjusted-*p* = .005), salience (*β* = -0.30, adjusted-*p* = .009), default mode (*β* = -0.31, adjusted-*p* < .001), reward (*β* = -0.55, adjusted-*p* = .001), dorsal somatomotor (*β* = -0.50, adjusted-*p* < .001), and visual (*β* = -0.38, adjusted-*p* < .001) networks. During the rewarded go/no-go task, we also observed also significantly increased within-module degree in the fronto-parietal network (*β* = 0.30, adjusted-*p* < .005). All other adjusted-*p*s > .203 (**Supplementary Table 3**).

### Differences in nodal properties of brain regions involved specifically in response inhibition, attention, and reward processing

Next, we investigated nodal properties of 25 brain regions derived from Neurosynth meta-analysis maps of the terms response inhibition, attention, or reward. For each of these brain regions, we quantified differences in node dissociation index and within module degree between pairs of cognitive states. During the standard go/no-go task as compared to the resting state, we observed significantly decreased node dissociation index in one attention-related node (*β* = -0.26, adjusted-*p* = .032) and three reward-related nodes (*β*s between -0.44 and -0.29, adjusted-*p*s between .002 and .032), as well as significantly increased node dissociation index in one node implicated in both response inhibition and reward (*β* = 0.39, adjusted-*p* = .032). We also observed significantly increased within module degree in one response inhibition-related node (*β* = 0.33, adjusted-*p* = .016), two attention-related nodes (*β*s between 0.33 and 0.34, adjusted-*p*s both equal to .016), and one reward-related node (*β* = 0.26, adjusted-*p* = .032), as well as decreased within module degree in one attention-related node (*β* = -0.26, adjusted-*p* = .032), one reward-related node (*β* = -0.24, adjusted-*p* = .040), and a node implicated in both response inhibition and reward (*β* = -0.26, adjusted-*p* = .027). All other adjusted-*p*s > .05 (**Supplementary Table 4**).

During the rewarded go/no-go task compared to the standard go/no-go task, we observed decreased node dissociation index in seven reward-related nodes (*β*s between -0.54 and -0.21, adjusted-*p*s between .005 and .032). Additionally, we observed increased within module degree in one response inhibition-related node (*β* = 0.28, adjusted-*p* = .005). All other adjusted-*p*s > .05 (**Supplementary Table 5**).

## Discussion

We investigated differences in task performance and functional brain network organization between the resting state, a standard go/no-go task, and a rewarded go/no-go task in children between the ages of 8 and 12 years on the whole-brain, network, and regional levels. Behaviorally, we observed that commission error rate (our index of response inhibition), but not tau (our index of sustained attention), improved during the rewarded go/no-go task compared to the standard go/no-go task. Additionally, in terms of global and network-level brain organization, our findings revealed that the standard go/no-go task evoked less integrated brain network organization (i.e., lower global efficiency, node dissociation index) and more segregated brain network organization (i.e., higher modularity, within-module degree) compared to the resting state, and that these differences were further enhanced in the presence of rewards. Our regional analyses similarly reflected these shifts to a less integrated and more segregated organization when focusing on the topological role of specific regions implicated in response inhibition, attention, and reward processing. These findings improve our understanding of how functional brain network organization across whole-brain, network, and regional levels supports response inhibition and sustained attention in the presence and absence of rewards in late childhood.

### Rewards shape go/no-go task performance

First, we observed that commission error rate significantly decreased from the standard to the rewarded go/no-go task. This is consistent with prior literature observing improvements in response inhibition performance with the addition of rewards (Geier and Luna 2009; Smith et al. 2011; Winter and Sheridan 2014; Demurie et al. 2016; Fortenbaugh et al. 2017; Miyasaka and Nomura 2019; Burton et al. 2021). We additionally observed that higher self-reported responsivity to rewards as indexed by the BIS/BAS scale was related to significantly greater decreases in commission error rate and a trend toward greater decreases in tau on the rewarded go/no-go task relative to the standard go/no-go task. These results suggest that rewards improve response inhibition task performance, although the impact of reward on sustained attention processes is not uniform across individuals and may be influenced by individual differences in responsivity to rewards.

Notably, commission error rate and tau were not significantly correlated during the standard go/no-go task. Thus, we conclude that the theorized processes probed by each, response inhibition and sustained attention respectively, can be relatively independently assessed. However, during the rewarded go/no-go task, greater inattention was significantly associated with more failures to correctly inhibit task responses, which suggests that rewards may unify the processes of response inhibition and sustained attention to a degree. Speculatively, it may be the case that in individuals who are more responsive to receiving rewards, sustained attention is enhanced to promote better response inhibition, thus strengthening the link between the two processes in the rewarded condition. In other words, we postulate that reward-driven brain network reconfiguration may serve to enhance the coordination of cognitive processes.

### Children’s brain network organization was segregated during the go/no-go task compared to rest

Our findings of decreased global efficiency and increased modularity during performance of a go/no-go task with and without rewards are consistent with prior literature observing increased segregation during sustained attention tasks in adults (Sadaghiani et al. 2015; Zuberer et al. 2021) and children (Barber et al. 2015; O’Halloran et al. 2018; Thomson et al. 2022). The modular network organization observed here and in studies of sustained attention has been theoretically and empirically suggested to facilitate strong local information processing within networks rather than global processing across many networks (Meunier et al. 2010; Sporns 2013; Bassett et al. 2015; Sadaghiani et al. 2015; Cohen and D’Esposito 2016). This pattern contrasts with studies examining response inhibition in adolescents and young adults (Mennes et al. 2012; Marek et al. 2015; Pan et al. 2020), as well as with studies examining other aspects of executive function, which have consistently reported increased integration as compared to the resting state in adults (Braun et al. 2015; Spielberg, Galarce, et al. 2015; Vatansever et al. 2015; Cohen and D’Esposito 2016; Schultz and Cole 2016; Shine et al. 2016; Hearne et al. 2017; Shine and Poldrack 2018; Rieck et al. 2021) and children (Le et al. 2020).

Our network-level analyses paralleled our whole-brain findings of decreased integration and increased segregation, as we observed decreases in node dissociation index of most individual brain networks and increased within-module degree of the fronto-parietal network. This suggests that the observed whole-brain shifts in organization are a result of widespread changes across many networks, rather than the result of changes in a select few networks. The only network that exhibited both within- and between-network changes was the fronto-parietal network, which has been theorized to support moment-to-moment processing (Dosenbach et al. 2006; Dosenbach et al. 2007; Dosenbach et al. 2008; Power and Petersen 2013). The fronto-parietal network is highly integrated in some contexts, as in working memory tasks (Cole et al. 2013; Shine et al. 2016), and less integrated in others, as in automatization of learning tasks (Mohr et al. 2016) and in our findings. Further, the fronto-parietal network exhibits high flexibility (i.e., reconfiguration between cognitive states) in both adults (Cole et al. 2013) and children (Le et al. 2020). Our results align with the notion supported by these two pieces of evidence that fronto-parietal reconfiguration is likely attuned to specific task demands.

In addition to reconfiguration of the fronto-parietal network and other ‘task-positive’ networks supporting externally-oriented functions, the default mode network, which supports internally-oriented processes such as episodic memory and theory of mind (Raichle 2015; Buckner and DiNicola 2019), also exhibited decreased integration during the standard go/no-go task relative to the resting state. Reduced integration of the default mode network is thought to reflect reduced interference of these internally-oriented processes into externally-oriented task processing and has shown benefits for task performance (Stevens et al. 2007; Spielberg, Miller, et al. 2015; Chung et al. 2020; Duffy et al. 2021).

The conformity of our results with prior literature on sustained attention (Barber et al. 2015; Sadaghiani et al. 2015; O’Halloran et al. 2018; Thomson et al. 2022), rather than response inhibition (Mennes et al. 2012; Marek et al. 2015; Pan et al. 2020), suggests that an attention-based cognitive strategy, rather than a response inhibition-based strategy, may have been employed in the performance of the go/no-go tasks in our participants. This is consistent with findings that performance on go/no-go tasks in children aged 4-12 years aligns with context monitoring, or non-inhibitory attentional control, rather than response inhibition processes (Winter and Sheridan 2014). These findings are further congruent with neuroimaging and electrophysiological evidence that context monitoring, rather than response suppression, is employed during response inhibition in young adults (Chatham et al. 2012). Together, this prior evidence supports the idea that an attention-based strategy explains the increasingly modular network organization observed during our go/no-go tasks. It is also important to note that cognitive strategies employed during executive function tasks, including response inhibition tasks, tend to shift across development (Munakata et al. 2012; Chevalier et al. 2014; Chevalier et al. 2015; Church et al. 2017; Zheng and Church 2021; Niebaum and Munakata 2023). Although in our sample of children between the ages of 8 and 12 years it is difficult to tease apart the influences of strategy and development, this possibility should be explored in samples with a wider age range and that are longitudinal.

It is worth mentioning that go/no-go tasks may probe several other cognitive processes in addition to response inhibition and sustained attention, including working memory (Criaud and Boulinguez 2013) and motor execution (Simmonds et al. 2008; Wessel 2018). We identified a pattern of organization during the go/no-go tasks that is distinct from the globally integrated organization exhibited during working memory in children (Le et al. 2020). As such, we think it’s unlikely that a working memory strategy was the predominant strategy during the go/no-go tasks in our participants. Our finding of increased modular organization during the go/no-go tasks accords with the pattern of reconfiguration observed during motor execution tasks probing rapid, automatic, and well-learned motor sequences (Bassett et al. 2015; Chen and Deem 2015; Cohen and D’Esposito 2016; Mohr et al. 2016). This may suggest that the go/no-tasks were so easy as to be automatic in our participants, resulting in a motor execution-based strategy. However, our behavioral results indicate that this is unlikely to be the case, as participants were not performing at ceiling (see commission error rates in **Table 4).** Additional work with tasks designed to distinguish across different cognitive processes and strategies is needed to further support our findings and interpretations. Finally, another possibility is that cognitive processes are engaged dynamically and transiently at too short of timescales to be detected with the methodology employed here. Evidence for this idea comes from prior studies in which we, and others, found that reduced integration may reflect an average of fluctuations between integrated and segregated configurations across time (Garrett et al. 2018; Fong et al. 2019; Duffy et al. 2021). As such, it is possible that switching between cognitive processes (and thus segregated/integrated configurations) throughout the go/no-go task may result in increased segregation and reduced integration assessed across minutes of the task.

### Children’s brain network organization was further segregated during the rewarded go/no-go task

Our finding that the addition of rewards to the go/no-go task evoked further reductions in global efficiency, increases in modularity, and reductions in network dissociation index across most networks is contrary to literature reporting that monetary rewards strengthen task-based functional connectivity between pairs of regions in distinct networks during cognitive control in adults (Dixon and Christoff 2012; Boehler et al. 2014; Teslovich et al. 2014; Bahlmann et al. 2015; Cubillo et al. 2019). It is possible that rewards push the brain into a task-adaptive state by strengthening task-specific signatures, which would be reflected in increased integration for many cognitive control tasks (Braun et al. 2015; Cohen and D’Esposito 2016; Shine et al. 2016; Le et al. 2020), but increased segregation during tasks probing sustained attention (Sadaghiani et al. 2015). This conclusion is supported by prior literature using multivariate pattern analysis (MVPA) and representational similarity analysis (RSA) to predict task-related features.

Specifically, it was reported that task state during a cued switching task was better predicted by fronto-parietal activity during rewarded trials than nonrewarded trials using MVPA (Etzel et al. 2016). Similarly, Rothlein and colleagues (2018) used RSA and found that rewards strengthened stimulus representations in the default mode network and dorsal attention network during a continuous performance task (Rothlein et al. 2018). Our study extends these findings to network functioning and suggests that the addition of rewards in the rewarded go/no-go task may have strengthened the optimal network organization (i.e., increased segregation and decreased integration) of the standard go/no-go task. Similar to the change from resting state to the standard go/no-go task, we observed that the decreased integration was largely distributed across most of the brain networks examined.

An alternative explanation for the discrepancy between prior literature and our results could lie in the methodology. Our study investigates distributed functional connectivity across the whole brain, rather than between specific pairs of regions. It is possible that specific connections may be strengthened in the presence of rewards, while the overall topology of the whole brain becomes further segregated.

### Response inhibition, attention, and reward-related regions reconfigure with standard and rewarded go/no-go task demands

While the above results characterize whole brain and network-level changes in organization between the three cognitive states, these changes may not apply uniformly to all task-relevant nodes. As such, we investigated integration (node dissociation index) and segregation (within-module degree) of key regions implicated in response inhibition, attention, and reward-processing, extracted from the Neurosynth meta-analytic database (Yarkoni et al. 2011). In general, we found decreased node dissociation index and increased within module degree during the standard go/no-go task compared to the resting state. This pattern was evident in nodes implicated in response inhibition (n=1), attention (n=2), and reward processing (n=3), that were included in the Seitzman atlas-defined fronto-parietal, default mode, and reward networks. As such, the reconfiguration of these nodes between rest and the standard go/no-go is consistent with the general whole-network findings discussed above.

Similarly, during the rewarded go/no-go task compared to the standard go/no-go task, we observed decreased node dissociation index and increased within module degree in several nodes implicated in reward (n=7) and one node implicated in both reward and response inhibition. Similar to the nodes changing their role between the resting state and the standard go/no-go task, these nodes were also included in the Seitzman atlas-defined fronto-parietal, default mode, and reward networks. Again, this pattern is consistent with the general whole-network patterns of reconfiguration. Additionally, the preponderance of change in reward-related regions over response inhibition- and attention-related regions suggests that the widespread network changes seen with the addition of rewards may be largely due to changes in reward-related regions distributed across several higher-order cognitive networks, rather than monetary rewards changing connectivity of regions specifically implicated in response inhibition and attention.

Three nodes exhibited a different reconfiguration pattern. A dorsal attention node implicated in attention and a ventral attention node implicated in reward both exhibited decreased within module degree during the standard go/no-go task relative to the resting state, which indicates a shift away from strong local processing. Additionally, a dorsal region of the posterior cingulate cortex that is a component of the parieto-medial network (Seitzman et al. 2020) and included in the Neurosynth maps for both response inhibition and reward exhibited increased node dissociation index and decreased within module degree during the standard go/no-go task relative to the resting state. This region is thought to function as a switch board that shapes internally- and externally-directed attention by coordinating anticorrelations between the default mode and task-positive networks, such as the fronto-parietal and dorsal attention networks (Leech and Sharp 2014). There is evidence that the posterior cingulate cortex adjusts connectivity patterns based on arousal level and the breadth of attentional focus (for a comprehensive review, see (Leech and Sharp 2014)). As such, increased integration of the posterior cingulate cortex during the standard go/no-go task relative to the resting state could reflect that coordination of this region with other networks is important for adapting brain network organization for the go/no-go task.

## Conclusion

In summary, in a sample of children aged 8 to 12 years, we observed that brain network integration decreased and segregation increased during a standard go/no-go task relative to the resting state, and that the addition of rewards further decreased integration and increased segregation both globally throughout the brain and within several key networks. Organization of individual brain regions thought to be related to response inhibition, attention, and reward processing exhibited largely the same pattern. Given the correspondence of these patterns with those involved in the sustained attention literature, we theorize that children employed an attention-based task strategy, rather than a response inhibition-based strategy, during the go/no-go tasks, and that this strategy was further engaged with the addition of rewards.

Although our results identify improvement in response inhibition task performance and shifts in brain network integration and segregation between the standard and rewarded go/no-go tasks, our sample size was limited, and future studies with larger samples are necessary to directly relate brain network organization to behavioral metrics of task performance in standard and rewarded versions of cognitive tasks. Additionally, it remains unknown how maturational trajectories of brain network organization are related to changes in sustained attention, as well as in response inhibition, across development. Longitudinal studies are needed to assess these relationships, as well as the shift in cognitive strategies, across childhood and adolescence. These findings will ultimately increase our understanding of response inhibition and sustained attention processes in children, which are essential for higher-order cognition and life outcomes.

## Data and Code Availability

Data and code used in this manuscript is available upon request to the senior author (JRC).

## Author Contributions

Mackenzie E Mitchell (Conceptualization, Formal analysis, Methodology, Visualization, Writing – original draft, Writing – review & editing), Teague R Henry (Methodology, Software, Writing – review & editing), Nicholas D Fogleman (Writing – review & editing), Cleanthis Michael (Data curation, Validation, Writing – review & editing), Tehila Nugiel (Validation, Writing – review & editing), Jessica R Cohen (Conceptualization, Funding acquisition, Methodology, Project administration, Resources, Supervision, Writing – review & editing)

## Acknowledgements

We would like to thank the Biomedical Research Imaging Center at UNC, as well as Cheyenne L. Bricken and Kelly H. Eom for their assistance with data collection and project management. Additionally, we would like to thank Alexa Monachino and the undergraduate research assistants who participated in data collection and management. Finally, we would like to thank our participating families for their time and effort.

