## Supplemental Materials for "Differential reconfiguration of brain networks in children in response to standard versus rewarded go/no-go task demands"

### Supplementary Methods

*fMRI data processing*

Results included in this manuscript come from preprocessing performed using fMRIPrep 1.5.0 (Esteban et al. 2018; Esteban et al. 2019; RRID:SCR_016216), which is based on Nipype 1.2.2 (Gorgolewski et al. 2011; Esteban et al. 2020; RRID:SCR_002502).

*Anatomical data preprocessing:* A total of 1 or 2 T1-weighted (T1w) images were found within each input BIDS dataset. All of them were corrected for intensity non-uniformity (INU) with N4BiasFieldCorrection (Tustison et al. 2010), distributed with ANTs 2.2.0 (Avants et al. 2008; RRID:SCR_004757). The T1w-reference was then skull-stripped with a Nipype implementation of the antsBrainExtraction.sh workflow (from ANTs), using OASIS30ANTs as target template. Brain tissue segmentation of cerebrospinal fluid (CSF), white-matter (WM) and gray-matter (GM) was performed on the brain-extracted T1w using fast (FSL 5.0.9; RRID:SCR_002823; Zhang et al. 2001). A T1w-reference map was computed after registration of 2 T1w images (after INU-correction) using mri_robust_template (FreeSurfer 6.0.1; Reuter et al. 2010). Brain surfaces were reconstructed using recon-all (FreeSurfer 6.0.1; RRID:SCR_001847; Dale et al. 1999), and the brain mask estimated previously was refined with a custom variation of the method to reconcile ANTs-derived and FreeSurfer-derived segmentations of the cortical gray-matter of Mindboggle (RRID:SCR_002438; Klein et al. 2017). Volume-based spatial normalization to one standard space (MNI152NLin2009cAsym) was performed through nonlinear registration with antsRegistration (ANTs 2.2.0), using brain-extracted versions of both the T1w reference and the T1w template. The following template was selected for spatial normalization: ICBM 152 Nonlinear Asymmetrical template version 2009c (Fonov et al. 2009; RRID:SCR_008796; TemplateFlow ID: MNI152NLin2009cAsym).

*Functional data preprocessing:* For each of the BOLD runs found per participant (across all tasks and sessions), the following preprocessing was performed. First, a reference volume and its skull-stripped version were generated using a custom methodology of fMRIPrep. A deformation field to correct for susceptibility distortions was estimated based on fMRIPrep's fieldmap-less approach. The deformation field is that resulting from co-registering the BOLD reference to the same-participant T1w-reference with its intensity inverted (Huntenburg 2014; Wang et al. 2017). Registration was performed with antsRegistration (ANTs 2.2.0), and the process regularized by constraining deformation to be nonzero only along the phase-encoding direction, and modulated with an average fieldmap template (Treiber et al. 2016). Based on the estimated susceptibility distortion, an unwarped BOLD reference was calculated for a more accurate co-registration with the anatomical reference. The BOLD reference was then co-registered to the T1w reference using bbregister (FreeSurfer), which implements boundary-based registration (Greve and Fischl 2009). Co-registration was configured with six degrees of freedom. Head-motion parameters with respect to the BOLD reference (transformation matrices, and six corresponding rotation and translation parameters) were estimated before any spatiotemporal filtering using mcflirt (FSL 5.0.9; Jenkinson et al. 2002). BOLD runs were slice-time corrected using 3dTshift from AFNI 20160207 (Cox and Hyde 1997; RRID:SCR_005927). The BOLD timeseries were resampled to surfaces on the following spaces: fsaverage5. The BOLD time-series (including slice-timing correction) were resampled onto their original, native space by applying a single, composite transform to correct for head-motion and susceptibility distortions. These resampled BOLD timeseries will be referred to as 'preprocessed BOLD in original space,' or just 'preprocessed BOLD.' The BOLD timeseries were resampled into standard space, generating a preprocessed BOLD run in ['MNI152NLin2009cAsym'] space. Confounding time-series were calculated based on the preprocessed BOLD framewise displacement (FD). FD was calculated for each functional run, using the implementation in Nipype (following the definition by Power et al. 2014). The head-motion estimates calculated in the correction step were also placed within the corresponding confounds file. The confound timeseries derived from head motion estimates were expanded with the inclusion of temporal derivatives and quadratic terms for each (Satterthwaite et al. 2013). All resamplings were performed with a single interpolation step by composing all the pertinent transformations (i.e. ~head-motion transform matrices, susceptibility distortion correction when available, and co-registrations to anatomical and output spaces). Gridded (volumetric) resamplings were performed using antsApplyTransforms (ANTs), configured with Lanczos interpolation to minimize the smoothing effects of other kernels (Lanczos, C 1964). Non-gridded (surface) resamplings were performed using mri_vol2surf (FreeSurfer).

Many internal operations of fMRIPrep use Nilearn 0.5.2 (Abraham et al. 2014; RRID:SCR_001362), mostly within the functional processing workflow. For more details of the pipeline, see the section corresponding to workflows in fMRIPrep's documentation.

*Copyright Waiver:* The above boilerplate text was automatically generated by fMRIPrep with the express intention that users should copy and paste this text into their manuscripts unchanged. Edits were only made to correct grammatical errors. It is released under the CC0 license.

*fMRI data quality checking*

To ensure that our fMRI data processing measures sufficiently removed the impacts of head motion on functional connectivity, we calculated the benchmarks reported by Ciric and colleagues (2017; **Supplementary Figure 1**). We found that the absolute median correlation between framewise displacement (FD) and functional connectivity strength (i.e., QC-FC correlation) was $r$ = 0.124, which would rank moderately among the methods tested on low motion data by Ciric and colleagues (2017). Additionally, the correlation between the Euclidean distance between pairs of regions and the QC-FC correlation of each pair (i.e., distance-dependent motion effects) was $r$ = -0.144, which would rank among the best methods tested by Ciric and colleagues (Ciric et al. 2017).

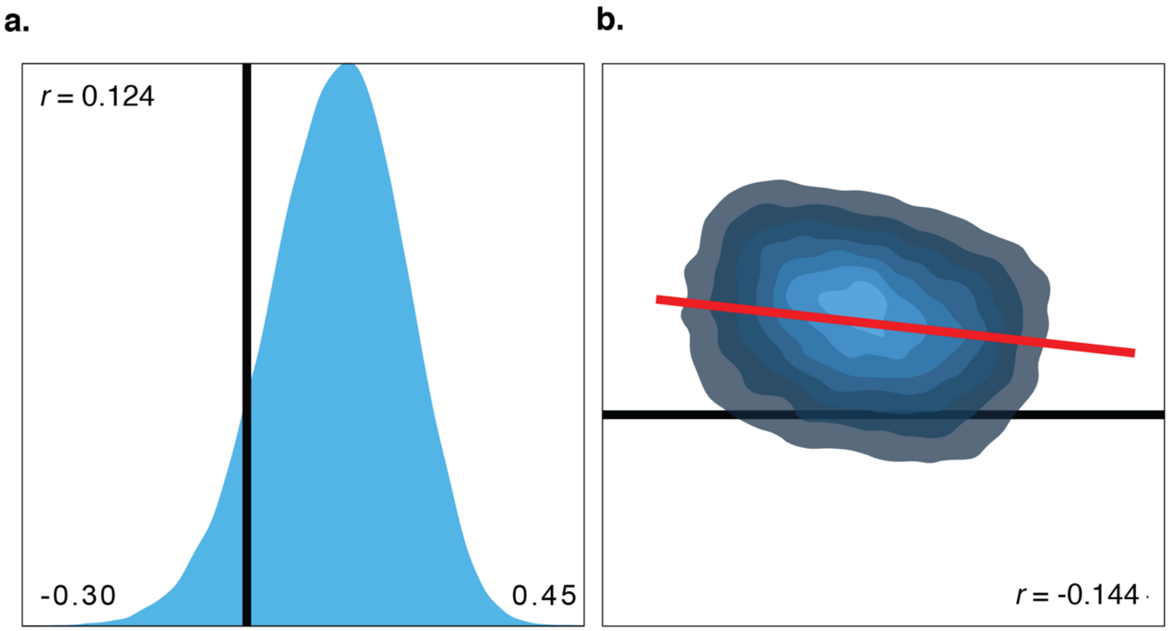


**Supplementary Figure 1. Relationships between residual effects of motion and functional connectivity estimates. a**. The distribution and absolute median correlation of QC-FC correlations between participant head motion (framewise displacement; FD) and functional connectivity strength. **b**. Density plot of distance-dependent motion effects where the x-axis is the Euclidean distance between pairs of regions and the y-axis is QC-FC correlation of each pair. Plots were generated via eXtensible Connectivity Pipeline (XCP) software (https://github.com/PennBBL/xcpEngine; Ciric et al. 2018).

*Graph thresholding*

To determine which graph thresholds to include in the final analyses, percent of disconnected nodes was calculated for each network of interest at each absolute threshold value (**Supplementary Figure 2**). The elbow point was calculated to determine the point at which thresholding induced a sharp increase in disconnected nodes. The thresholds below the elbow point were retained for the analyses (*r* = 0.20 – 0.30).

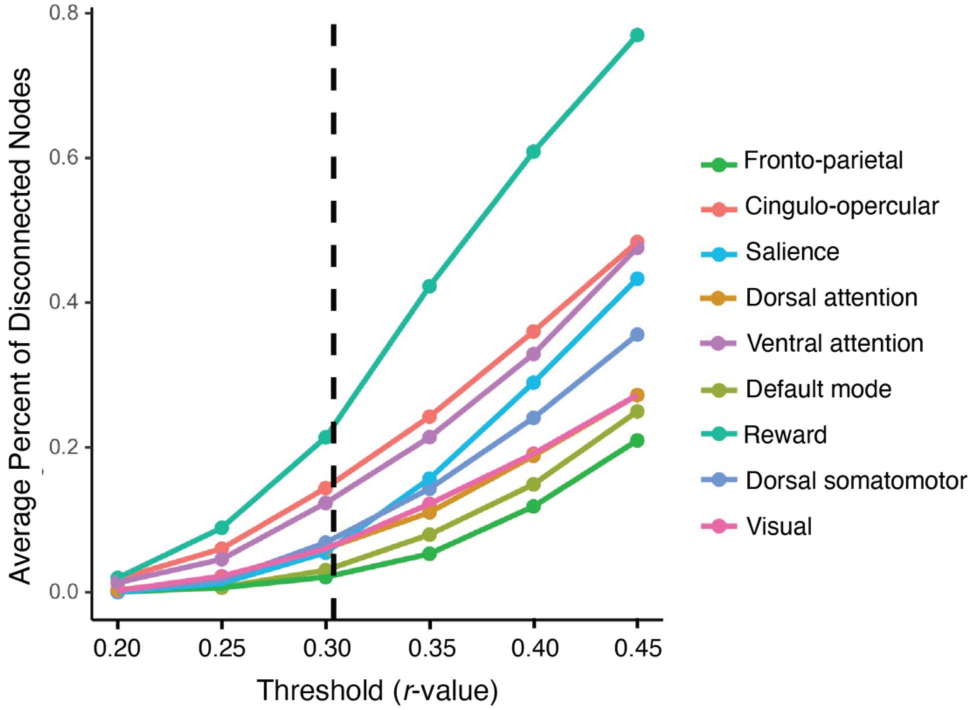


**Supplementary Figure 2. Determination of the elbow point cutoff for graph thresholding.** The percent of disconnected nodes for each network of interest averaged across all participants and cognitive states and shown for each absolute threshold. The elbow point (vertical black dashed line) was determined to be between the 0.30 and 0.35 thresholds.

### Supplementary Results

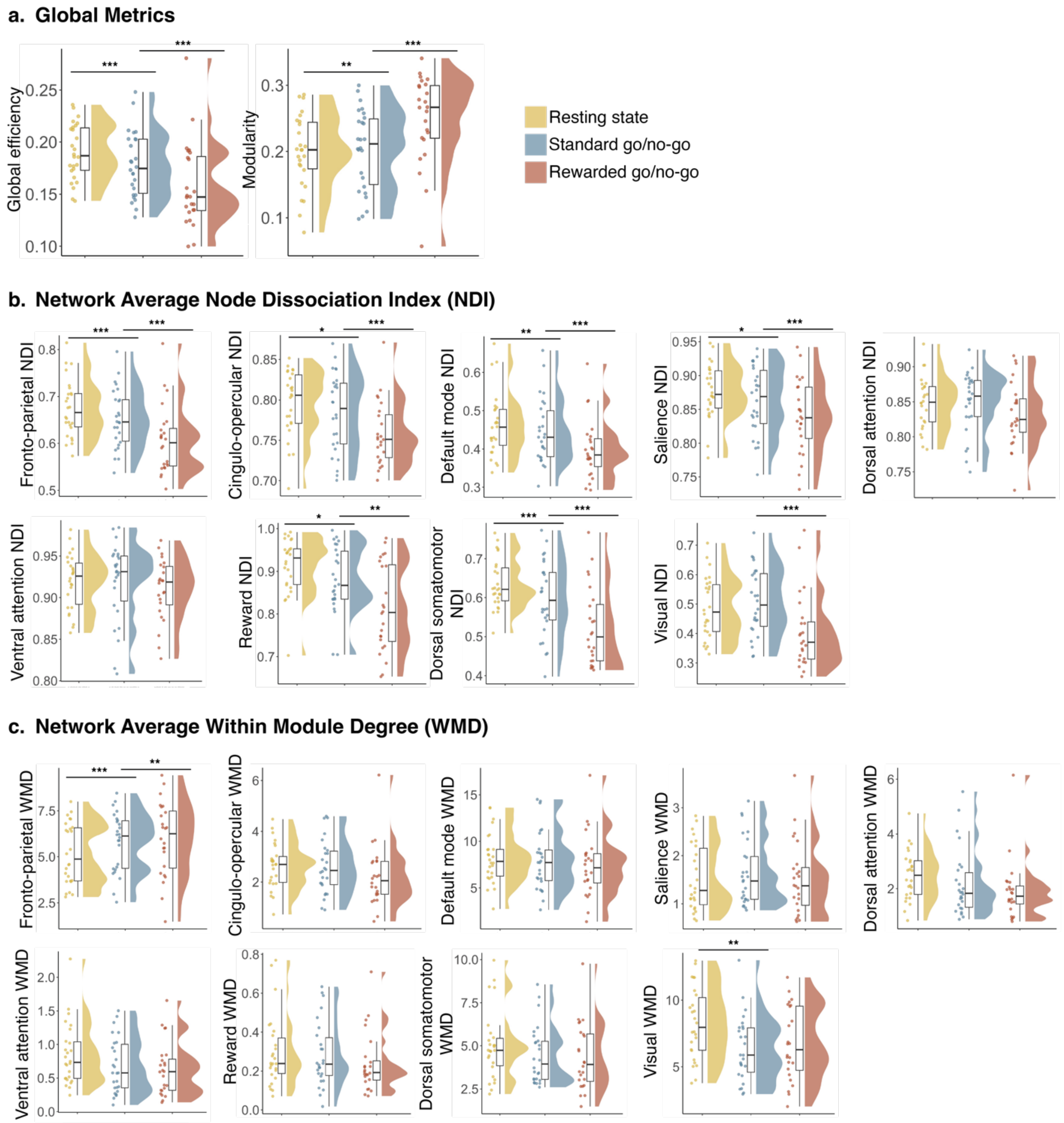


**Figure 3. The distribution of each graph metric in the resting state, standard go/no-go task, and rewarded go/no-go task. a**. The distribution of global metrics (global efficiency, modularity). **b**. The distribution of average node dissociation index (NDI) for each network of interest. **b**. The distribution of average within module degree (WMD) for each network of interest. *adjusted-*p* < .05, **adjusted-*p* < .01, ***adjusted-*p* < .001

**Supplementary Table 1.** Differences in whole brain organization between the resting state, standard go/no-go task, and rewarded go/no-go task.

| *Brain metric* | *Comparison* | *b* | *β* | *SE* | *t* | *df* | *p* | *adjusted-p* |
|---|---|---|---|---|---|---|---|---|
| Global efficiency | Resting state vs. standard go/no-go task | -0.01 | -0.26 | 0.001 | -7.82 | 49.90 | 3.16E-10 | **6.32E-10** |
| Global efficiency | Standard go/no-go task vs. rewarded go/no-go task | -0.01 | -0.32 | 0.001 | -9.34 | 49.77 | 1.59E-12 | **3.18E-12** |
| Modularity | Resting state vs. standard go/no-go task | 0.01 | 0.17 | 0.004 | 2.78 | 47.48 | 0.008 | **0.008** |
| Modularity | Standard go/no-go task vs. rewarded go/no-go task | 0.02 | 0.36 | 0.004 | 5.70 | 47.27 | 7.46E-07 | **7.46E-07** |

Standard errors are for the unstandardized betas. All models include graph density, graph mean FC, age, and sex as covariates of no interest. $\beta$ is the standardized beta coefficient. Standard errors are for the unstandardized betas. Significant adjusted-*p* values < .05 are denoted in bold.

**Supplementary Table 2.** Differences in network organization between the resting state and the standard go/no-go task.

| *Brain metric* | *b* | *β* | *SE* | *t* | *df* | *p* | *adjusted-p* |
|---|---|---|---|---|---|---|---|
| NDI of the fronto-parietal network | -0.02 | -0.29 | 0.004 | -4.60 | 47.22 | 3.20E-05 | **0.0003** |
| NDI of the cingulo-opercular network | -0.01 | -0.26 | 0.005 | -2.49 | 49.80 | 0.016 | **0.036** |
| NDI of the default mode network | -0.02 | -0.23 | 0.005 | -3.69 | 45.50 | 0.001 | **0.003** |
| NDI of the salience network | -0.01 | -0.25 | 0.005 | -2.60 | 49.94 | 0.012 | **0.032** |
| NDI of the reward network | -0.04 | -0.44 | 0.013 | -3.06 | 48.99 | 0.004 | **0.011** |
| NDI of the dorsal somatomotor network | -0.04 | -0.42 | 0.007 | -5.68 | 48.45 | 7.55E-07 | **1.36E-05** |
| NDI of the visual network | -0.01 | -0.08 | 0.009 | -1.05 | 49.80 | 0.297 | 0.411 |
| NDI of the dorsal attention network | 0.00 | -0.02 | 0.004 | -0.23 | 48.26 | 0.823 | 0.871 |
| NDI of the ventral attention network | 0.00 | -0.05 | 0.005 | -0.36 | 50.53 | 0.724 | 0.814 |
| WMD of the fronto-parietal network | 0.67 | 0.38 | 0.157 | 4.25 | 46.78 | 0.0001 | **0.001** |
| WMD of the cingulo-opercular network | 0.03 | 0.03 | 0.068 | 0.43 | 50.06 | 0.669 | 0.814 |
| WMD of the default mode network | -0.02 | -0.01 | 0.194 | -0.11 | 44.18 | 0.912 | 0.912 |
| WMD of the salience network | 0.11 | 0.16 | 0.055 | 2.04 | 48.40 | 0.047 | 0.084 |
| WMD of the reward network | -0.01 | -0.04 | 0.020 | -0.37 | 45.69 | 0.710 | 0.814 |
| WMD of the dorsal somatomotor network | -0.15 | -0.08 | 0.129 | -1.19 | 48.68 | 0.238 | 0.357 |
| WMD of the visual network | -0.76 | -0.28 | 0.225 | -3.37 | 49.59 | 0.001 | **0.005** |
| WMD of the dorsal attention network | -0.15 | -0.14 | 0.090 | -1.69 | 49.47 | 0.097 | 0.159 |

| | | | | | | | |
|---|---|---|---|---|---|---|---|
| WMD of the ventral attention network | -0.09 | -0.20 | 0.038 | -2.23 | 49.54 | 0.031 | 0.061 |

Standard errors are for the unstandardized betas. All models include graph density, graph mean FC, age, and sex as covariates of no interest. $\beta$ is the standardized beta coefficient. Standard errors are for the unstandardized betas. NDI = node dissociation index. WMD = within-module degree. Significant adjusted-*p* values < .05 are denoted in bold.

**Supplementary Table 3.** Differences in network organization between the standard go/no-go task and the rewarded go/no-go task.

| ***Brain metric*** | ***b*** | $\beta$ | ***SE*** | ***t*** | ***df*** | ***p*** | ***adjusted-p*** |
|---|---|---|---|---|---|---|---|
| NDI of the fronto-parietal network | -0.03 | -0.39 | 0.005 | -6.07 | 46.99 | 2.10E-07 | **1.89E-06** |
| NDI of the cingulo-opercular network | -0.02 | -0.34 | 0.005 | -3.25 | 49.89 | 0.002 | **0.005** |
| NDI of the default mode network | -0.03 | -0.31 | 0.006 | -4.86 | 45.28 | 1.44E-05 | **6.50E-05** |
| NDI of the salience network | -0.01 | -0.30 | 0.005 | -3.02 | 49.79 | 0.004 | **0.009** |
| NDI of the reward network | -0.05 | -0.55 | 0.013 | -3.82 | 49.55 | 0.0004 | **0.001** |
| NDI of the dorsal somatomotor network | -0.05 | -0.50 | 0.007 | -6.70 | 48.43 | 2.07E-08 | **3.72E-07** |
| NDI of the visual network | -0.05 | -0.38 | 0.009 | -4.93 | 49.60 | 9.71E-06 | **5.82E-05** |
| NDI of the dorsal attention network | -0.01 | -0.17 | 0.004 | -1.67 | 48.09 | 0.102 | 0.203 |
| NDI of the ventral attention network | 0.00 | -0.11 | 0.005 | -0.83 | 50.57 | 0.411 | 0.528 |
| WMD of the fronto-parietal network | 0.53 | 0.30 | 0.160 | 3.29 | 46.55 | 0.002 | **0.005** |
| WMD of the cingulo-opercular network | -0.06 | -0.06 | 0.069 | -0.91 | 49.89 | 0.368 | 0.510 |
| WMD of the default mode network | -0.13 | -0.04 | 0.198 | -0.63 | 43.94 | 0.530 | 0.597 |
| WMD of the salience network | 0.05 | 0.07 | 0.056 | 0.92 | 48.17 | 0.361 | 0.510 |
| WMD of the reward network | -0.01 | -0.07 | 0.020 | -0.63 | 45.81 | 0.531 | 0.597 |
| WMD of the dorsal somatomotor network | 0.01 | 0.01 | 0.132 | 0.11 | 48.45 | 0.914 | 0.966 |
| WMD of the visual network | -0.01 | 0.00 | 0.229 | -0.04 | 49.38 | 0.966 | 0.966 |
| WMD of the dorsal attention network | -0.13 | -0.12 | 0.092 | -1.41 | 49.30 | 0.164 | 0.295 |
| WMD of the ventral attention network | -0.04 | -0.08 | 0.039 | -0.92 | 49.32 | 0.360 | 0.510 |

Standard errors are for the unstandardized betas. All models include graph density, graph mean FC, age, and sex as covariates of no interest. $\beta$ is the standardized beta coefficient. Standard errors are for the unstandardized betas. NDI = node dissociation index. WMD = within-module degree. Significant adjusted-*p* values < .05 are denoted in bold.

**Supplementary Table 4.** Differences in nodal organization between the resting state and the standard go/no-go task.

| *Brain metric* | *ROI* | *X* | *Y* | *Z* | *Neurosynth map* | *Seitzman network* | *b* | *β* | *SE* | *t* | *df* | *p* | *adjusted-p* |
|---|---|---|---|---|---|---|---|---|---|---|---|---|---|
| NDI | 198 | 49.18 | -42.41 | 45.16 | Response inhibition | Fronto-parietal | -0.02 | -0.18 | 0.01 | -1.83 | 48.49 | 0.073 | 0.182 |
| NDI | 201 | 55.27 | -44.59 | 36.7 | Response inhibition | Fronto-parietal | -0.02 | -0.15 | 0.01 | -1.78 | 48.00 | 0.081 | 0.187 |
| NDI | 209 | 26.07 | 49.56 | 26.58 | Response inhibition | Salience | 0.00 | -0.05 | 0.01 | -0.49 | 49.59 | 0.627 | 0.799 |
| NDI | 210 | 33.56 | 16.45 | -7.58 | Response inhibition & Reward | Salience | -0.01 | -0.04 | 0.02 | -0.28 | 51.19 | 0.783 | 0.890 |
| NDI | 139 | 1.75 | -24.25 | 30.36 | Response inhibition & Reward | Parieto-medial | 0.02 | 0.39 | 0.01 | 2.81 | 50.63 | 0.007 | **0.032** |
| NDI | 184 | -28.4 | -57.93 | 47.78 | Attention | Fronto-parietal | -0.04 | -0.26 | 0.01 | -2.89 | 45.63 | 0.006 | **0.032** |
| NDI | 189 | 33.38 | -53.12 | 44.02 | Attention | Fronto-parietal | -0.02 | -0.14 | 0.01 | -1.78 | 45.37 | 0.082 | 0.187 |
| NDI | 229 | 9.61 | -61.5 | 60.88 | Attention | Dorsal attention | 0.01 | 0.06 | 0.01 | 0.48 | 49.91 | 0.630 | 0.799 |
| NDI | 230 | 21.9 | -64.74 | 48.12 | Attention | Dorsal attention | 0.00 | 0.02 | 0.01 | 0.23 | 48.15 | 0.816 | 0.907 |
| NDI | 231 | 25.34 | -58.18 | 60.34 | Attention | Dorsal attention | 0.00 | -0.01 | 0.01 | -0.10 | 50.34 | 0.924 | 0.978 |
| NDI | 232 | 28.56 | -4.62 | 53.99 | Attention | Dorsal attention | 0.01 | 0.07 | 0.01 | 0.73 | 50.02 | 0.468 | 0.684 |
| NDI | 250 | 14.15 | -1.19 | 18.18 | Reward | Fronto-parietal | -0.03 | -0.25 | 0.01 | -2.29 | 50.50 | 0.026 | 0.088 |
| NDI | 248 | 11.87 | 17.51 | 6.57 | Reward | Salience | -0.01 | -0.11 | 0.02 | -0.71 | 50.67 | 0.479 | 0.684 |
| NDI | 249 | -13.28 | 17.24 | 7.14 | Reward | Salience | -0.01 | -0.13 | 0.01 | -0.89 | 50.71 | 0.377 | 0.589 |
| NDI | 108 | -3.06 | 44.41 | -9.46 | Reward | Default mode | -0.05 | -0.29 | 0.02 | -2.92 | 45.89 | 0.005 | **0.032** |
| NDI | 110 | -2.5 | 41.7 | 16.05 | Reward | Default mode | 0.00 | 0.01 | 0.02 | 0.10 | 48.98 | 0.920 | 0.978 |
| NDI | 116 | 7.51 | 42.49 | -5.35 | Reward | Default mode | -0.06 | -0.35 | 0.01 | -4.51 | 48.10 | 0.000 | **0.002** |
| NDI | 117 | 8.36 | 47.59 | -15.18 | Reward | Default mode | -0.04 | -0.29 | 0.02 | -2.56 | 45.76 | 0.014 | 0.053 |
| NDI | 262 | 3.42 | -7.79 | 8.23 | Reward | Default mode | -0.02 | -0.08 | 0.02 | -1.04 | 47.80 | 0.301 | 0.523 |
| NDI | 256 | 25.26 | 1.75 | -1.26 | Reward | Ventral attention | 0.00 | 0.05 | 0.01 | 0.32 | 69.00 | 0.752 | 0.874 |

| | | | | | | | | | | | | |
|---|---|---|---|---|---|---|---|---|---|---|---|---|
| NDI | 241 | -21.14 | 40.87 | -20.48 | Reward | Reward | 0.00 | -0.10 | 0.00 | -0.60 | 69.00 | 0.550 | 0.744 |
| NDI | 242 | 23.96 | 31.94 | -17.78 | Reward | Reward | 0.01 | 0.20 | 0.00 | 1.29 | 50.73 | 0.204 | 0.424 |
| NDI | 245 | 19.51 | -1.85 | -23.11 | Reward | Reward | -0.07 | -0.30 | 0.03 | -2.10 | 43.21 | 0.042 | 0.123 |
| NDI | 246 | 12.66 | 17.32 | -5.06 | Reward | Reward | -0.06 | -0.31 | 0.03 | -2.42 | 48.23 | 0.019 | 0.069 |
| NDI | 247 | -12.49 | 17.05 | -4.49 | Reward | Reward | -0.11 | -0.44 | 0.03 | -3.30 | 69.00 | 0.002 | **0.016** |
| WMD | 198 | 49.18 | -42.41 | 45.16 | Response inhibition | Fronto-parietal | 1.04 | 0.33 | 0.29 | 3.54 | 45.23 | 0.001 | **0.016** |
| WMD | 201 | 55.27 | -44.59 | 36.7 | Response inhibition | Fronto-parietal | 0.59 | 0.19 | 0.29 | 1.99 | 48.22 | 0.052 | 0.146 |
| WMD | 209 | 26.07 | 49.56 | 26.58 | Response inhibition | Salience | 0.20 | 0.21 | 0.10 | 2.09 | 49.68 | 0.041 | 0.123 |
| WMD | 210 | 33.56 | 16.45 | -7.58 | Response inhibition & Reward | Salience | 0.08 | 0.09 | 0.09 | 0.90 | 49.53 | 0.374 | 0.589 |
| WMD | 139 | 1.75 | -24.25 | 30.36 | Response inhibition & Reward | Parieto-medial | -0.16 | -0.26 | 0.05 | -3.10 | 48.00 | 0.003 | **0.027** |
| WMD | 184 | -28.4 | -57.93 | 47.78 | Attention | Fronto-parietal | 1.03 | 0.33 | 0.29 | 3.60 | 46.88 | 0.001 | **0.016** |
| WMD | 189 | 33.38 | -53.12 | 44.02 | Attention | Fronto-parietal | 1.13 | 0.34 | 0.33 | 3.43 | 47.34 | 0.001 | **0.016** |
| WMD | 229 | 9.61 | -61.5 | 60.88 | Attention | Dorsal attention | -0.14 | -0.10 | 0.14 | -1.06 | 49.62 | 0.296 | 0.523 |
| WMD | 230 | 21.9 | -64.74 | 48.12 | Attention | Dorsal attention | 0.00 | 0.00 | 0.15 | -0.02 | 48.83 | 0.987 | 0.987 |
| WMD | 231 | 25.34 | -58.18 | 60.34 | Attention | Dorsal attention | -0.16 | -0.09 | 0.16 | -1.01 | 48.54 | 0.316 | 0.526 |
| WMD | 232 | 28.56 | -4.62 | 53.99 | Attention | Dorsal attention | -0.44 | -0.26 | 0.15 | -2.90 | 49.97 | 0.005 | **0.032** |
| WMD | 250 | 14.15 | -1.19 | 18.18 | Reward | Fronto-parietal | 0.19 | 0.06 | 0.26 | 0.74 | 47.93 | 0.464 | 0.684 |
| WMD | 248 | 11.87 | 17.51 | 6.57 | Reward | Salience | 0.09 | 0.10 | 0.08 | 1.11 | 46.73 | 0.272 | 0.523 |
| WMD | 249 | -13.28 | 17.24 | 7.14 | Reward | Salience | 0.00 | 0.00 | 0.08 | 0.02 | 48.04 | 0.982 | 0.987 |
| WMD | 108 | -3.06 | 44.41 | -9.46 | Reward | Default mode | 1.40 | 0.26 | 0.49 | 2.85 | 47.55 | 0.006 | **0.032** |
| WMD | 110 | -2.5 | 41.7 | 16.05 | Reward | Default mode | -0.39 | -0.10 | 0.21 | -1.84 | 46.55 | 0.072 | 0.182 |
| WMD | 116 | 7.51 | 42.49 | -5.35 | Reward | Default mode | 0.03 | 0.01 | 0.38 | 0.08 | 48.84 | 0.939 | 0.978 |
| WMD | 117 | 8.36 | 47.59 | -15.18 | Reward | Default mode | 0.47 | 0.11 | 0.36 | 1.29 | 48.97 | 0.202 | 0.424 |
| WMD | 262 | 3.42 | -7.79 | 8.23 | Reward | Default mode | -0.06 | -0.04 | 0.16 | -0.38 | 47.56 | 0.707 | 0.862 |

| | | | | | | | | | | | | | |
|---|---|---|---|---|---|---|---|---|---|---|---|---|---|
| WMD | 256 | 25.26 | 1.75 | -1.26 | Reward | Ventral attention | -0.11 | -0.24 | 0.04 | -2.69 | 49.17 | 0.010 | **0.040** |
| WMD | 241 | -21.14 | 40.87 | -20.48 | Reward | Reward | -0.02 | -0.11 | 0.03 | -0.68 | 69.00 | 0.500 | 0.695 |
| WMD | 242 | 23.96 | 31.94 | -17.78 | Reward | Reward | -0.02 | -0.05 | 0.04 | -0.47 | 47.40 | 0.639 | 0.799 |
| WMD | 245 | 19.51 | -1.85 | -23.11 | Reward | Reward | -0.01 | -0.05 | 0.03 | -0.32 | 40.55 | 0.750 | 0.874 |
| WMD | 246 | 12.66 | 17.32 | -5.06 | Reward | Reward | 0.04 | 0.15 | 0.03 | 1.13 | 46.41 | 0.264 | 0.523 |
| WMD | 247 | -12.49 | 17.05 | -4.49 | Reward | Reward | 0.04 | 0.15 | 0.04 | 1.04 | 42.73 | 0.303 | 0.523 |

Standard errors are for the unstandardized betas. All models include graph density, graph mean FC, age, and sex as covariates of no interest. $\beta$ is the standardized beta coefficient. Standard errors are for the unstandardized betas. NDI = node dissociation index. WMD = within-module degree. Significant adjusted-*p* values < .05 are denoted in bold. ROI number corresponds to the order of each region in the Seitzman atlas. Coordinates are reported in MNI space.

**Supplementary Table 5.** Differences in nodal organization between the standard go/no-go task and the rewarded go/no-go task.

| *Brain metric* | *ROI* | *X* | *Y* | *Z* | *Neurosynth map* | *Seitzman network* | *b* | $\beta$ | *SE* | *t* | *df* | *p* | *adjusted -p* |
|---|---|---|---|---|---|---|---|---|---|---|---|---|---|
| NDI | 198 | 49.18 | -42.41 | 45.16 | Response inhibition | Fronto-parietal | -0.02 | -0.19 | 0.01 | -1.93 | 48.27 | 0.059 | 0.197 |
| NDI | 201 | 55.27 | -44.59 | 36.7 | Response inhibition | Fronto-parietal | -0.02 | -0.17 | 0.01 | -2.05 | 47.80 | 0.046 | 0.186 |
| NDI | 209 | 26.07 | 49.56 | 26.58 | Response inhibition | Salience | -0.01 | -0.20 | 0.01 | -1.76 | 49.39 | 0.085 | 0.267 |
| NDI | 210 | 33.56 | 16.45 | -7.58 | Response inhibition & Reward | Salience | 0.00 | 0.04 | 0.02 | 0.23 | 51.55 | 0.822 | 0.928 |
| NDI | 139 | 1.75 | -24.25 | 30.36 | Response inhibition & Reward | Parieto-medial | 0.01 | 0.16 | 0.01 | 1.16 | 50.78 | 0.252 | 0.489 |
| NDI | 184 | -28.4 | -57.93 | 47.78 | Attention | Fronto-parietal | -0.02 | -0.13 | 0.02 | -1.48 | 45.38 | 0.146 | 0.349 |
| NDI | 189 | 33.38 | -53.12 | 44.02 | Attention | Fronto-parietal | -0.01 | -0.08 | 0.01 | -1.03 | 45.14 | 0.308 | 0.550 |
| NDI | 229 | 9.61 | -61.5 | 60.88 | Attention | Dorsal attention | 0.00 | -0.01 | 0.01 | -0.12 | 49.82 | 0.905 | 0.943 |
| NDI | 230 | 21.9 | -64.74 | 48.12 | Attention | Dorsal attention | -0.01 | -0.13 | 0.01 | -1.24 | 47.92 | 0.221 | 0.489 |
| NDI | 231 | 25.34 | -58.18 | 60.34 | Attention | Dorsal attention | -0.02 | -0.26 | 0.01 | -2.41 | 50.22 | 0.020 | 0.098 |

| | | | | | | | | | | | | | |
|---|---|---|---|---|---|---|---|---|---|---|---|---|---|
| NDI | 232 | 28.56 | -4.62 | 53.99 | Attention | Dorsal attention | -0.01 | -0.07 | 0.01 | -0.79 | 49.83 | 0.433 | 0.648 |
| NDI | 250 | 14.15 | -1.19 | 18.18 | Reward | Fronto-parietal | -0.04 | -0.32 | 0.01 | -2.94 | 50.40 | 0.005 | **0.032** |
| NDI | 248 | 11.87 | 17.51 | 6.57 | Reward | Salience | -0.03 | -0.27 | 0.02 | -1.71 | 51.14 | 0.093 | 0.272 |
| NDI | 249 | -13.28 | 17.24 | 7.14 | Reward | Salience | 0.01 | 0.09 | 0.01 | 0.64 | 50.90 | 0.524 | 0.689 |
| NDI | 108 | -3.06 | 44.41 | -9.46 | Reward | Default mode | -0.06 | -0.33 | 0.02 | -3.29 | 45.64 | 0.002 | **0.024** |
| NDI | 110 | -2.5 | 41.7 | 16.05 | Reward | Default mode | 0.00 | 0.02 | 0.02 | 0.16 | 48.77 | 0.871 | 0.943 |
| NDI | 116 | 7.51 | 42.49 | -5.35 | Reward | Default mode | -0.06 | -0.32 | 0.01 | -4.06 | 47.92 | 0.000 | **0.005** |
| NDI | 117 | 8.36 | 47.59 | -15.18 | Reward | Default mode | -0.05 | -0.31 | 0.02 | -2.67 | 45.92 | 0.010 | 0.058 |
| NDI | 262 | 3.42 | -7.79 | 8.23 | Reward | Default mode | -0.07 | -0.21 | 0.02 | -2.93 | 47.63 | 0.005 | **0.032** |
| NDI | 256 | 25.26 | 1.75 | -1.26 | Reward | Ventral attention | -0.01 | -0.12 | 0.01 | -0.74 | 69.00 | 0.462 | 0.648 |
| NDI | 241 | -21.14 | 40.87 | -20.48 | Reward | Reward | 0.01 | 0.32 | 0.00 | 1.98 | 69.00 | 0.052 | 0.186 |
| NDI | 242 | 23.96 | 31.94 | -17.78 | Reward | Reward | 0.01 | 0.19 | 0.00 | 1.21 | 51.09 | 0.232 | 0.489 |
| NDI | 245 | 19.51 | -1.85 | -23.11 | Reward | Reward | -0.11 | -0.48 | 0.03 | -3.20 | 43.54 | 0.003 | **0.025** |
| NDI | 246 | 12.66 | 17.32 | -5.06 | Reward | Reward | -0.09 | -0.44 | 0.03 | -3.33 | 48.27 | 0.002 | **0.024** |
| NDI | 247 | -12.49 | 17.05 | -4.49 | Reward | Reward | -0.13 | -0.54 | 0.03 | -3.92 | 69.00 | 0.000 | **0.005** |
| WMD | 198 | 49.18 | -42.41 | 45.16 | Response inhibition | Fronto-parietal | 0.88 | 0.28 | 0.30 | 2.94 | 44.99 | 0.005 | **0.032** |
| WMD | 201 | 55.27 | -44.59 | 36.7 | Response inhibition | Fronto-parietal | 0.21 | 0.07 | 0.30 | 0.71 | 47.99 | 0.479 | 0.648 |
| WMD | 209 | 26.07 | 49.56 | 26.58 | Response inhibition | Salience | 0.12 | 0.12 | 0.10 | 1.19 | 49.49 | 0.240 | 0.489 |
| WMD | 210 | 33.56 | 16.45 | -7.58 | Response inhibition & Reward | Salience | 0.08 | 0.10 | 0.09 | 0.90 | 49.38 | 0.372 | 0.620 |
| WMD | 139 | 1.75 | -24.25 | 30.36 | Response inhibition & Reward | Parieto-medial | -0.01 | -0.02 | 0.05 | -0.21 | 47.79 | 0.836 | 0.928 |
| WMD | 184 | -28.4 | -57.93 | 47.78 | Attention | Fronto-parietal | 0.22 | 0.07 | 0.29 | 0.75 | 46.66 | 0.455 | 0.648 |
| WMD | 189 | 33.38 | -53.12 | 44.02 | Attention | Fronto-parietal | 0.53 | 0.16 | 0.33 | 1.60 | 47.11 | 0.117 | 0.293 |

| | | | | | | | | | | | | | |
|---|---|---|---|---|---|---|---|---|---|---|---|---|---|
| WMD | 229 | 9.61 | -61.5 | 60.88 | Attention | Dorsal attention | -0.10 | -0.07 | 0.14 | -0.72 | 49.45 | 0.476 | 0.648 |
| WMD | 230 | 21.9 | -64.74 | 48.12 | Attention | Dorsal attention | -0.08 | -0.05 | 0.15 | -0.51 | 48.61 | 0.614 | 0.767 |
| WMD | 231 | 25.34 | -58.18 | 60.34 | Attention | Dorsal attention | -0.15 | -0.09 | 0.16 | -0.94 | 48.31 | 0.351 | 0.605 |
| WMD | 232 | 28.56 | -4.62 | 53.99 | Attention | Dorsal attention | -0.31 | -0.19 | 0.15 | -2.04 | 49.77 | 0.046 | 0.186 |
| WMD | 250 | 14.15 | -1.19 | 18.18 | Reward | Fronto-parietal | 0.43 | 0.14 | 0.26 | 1.61 | 47.70 | 0.113 | 0.293 |
| WMD | 248 | 11.87 | 17.51 | 6.57 | Reward | Salience | 0.14 | 0.16 | 0.08 | 1.68 | 46.58 | 0.099 | 0.275 |
| WMD | 249 | -13.28 | 17.24 | 7.14 | Reward | Salience | -0.02 | -0.02 | 0.08 | -0.22 | 48.01 | 0.827 | 0.928 |
| WMD | 108 | -3.06 | 44.41 | -9.46 | Reward | Default mode | 0.26 | 0.05 | 0.50 | 0.52 | 47.32 | 0.604 | 0.767 |
| WMD | 110 | -2.5 | 41.7 | 16.05 | Reward | Default mode | 0.03 | 0.01 | 0.22 | 0.12 | 46.44 | 0.903 | 0.943 |
| WMD | 116 | 7.51 | 42.49 | -5.35 | Reward | Default mode | -0.28 | -0.07 | 0.39 | -0.73 | 48.63 | 0.471 | 0.648 |
| WMD | 117 | 8.36 | 47.59 | -15.18 | Reward | Default mode | -0.02 | 0.00 | 0.37 | -0.05 | 48.75 | 0.963 | 0.963 |
| WMD | 262 | 3.42 | -7.79 | 8.23 | Reward | Default mode | 0.01 | 0.01 | 0.16 | 0.08 | 47.36 | 0.936 | 0.955 |
| WMD | 256 | 25.26 | 1.75 | -1.26 | Reward | Ventral attention | -0.05 | -0.10 | 0.04 | -1.15 | 49.15 | 0.255 | 0.489 |
| WMD | 241 | -21.14 | 40.87 | -20.48 | Reward | Reward | -0.05 | -0.32 | 0.03 | -1.98 | 69.00 | 0.052 | 0.186 |
| WMD | 242 | 23.96 | 31.94 | -17.78 | Reward | Reward | -0.03 | -0.09 | 0.04 | -0.79 | 47.79 | 0.436 | 0.648 |
| WMD | 245 | 19.51 | -1.85 | -23.11 | Reward | Reward | -0.01 | -0.07 | 0.03 | -0.45 | 40.65 | 0.658 | 0.802 |
| WMD | 246 | 12.66 | 17.32 | -5.06 | Reward | Reward | 0.04 | 0.15 | 0.03 | 1.12 | 46.41 | 0.267 | 0.494 |
| WMD | 247 | -12.49 | 17.05 | -4.49 | Reward | Reward | 0.01 | 0.03 | 0.04 | 0.22 | 43.07 | 0.824 | 0.928 |

Standard errors are for the unstandardized betas. All models include graph density, graph mean FC, age, and sex as covariates of no interest. $\beta$ is the standardized beta coefficient. Standard errors are for the unstandardized betas. NDI = node dissociation index. WMD = within-module degree. Significant adjusted-*p* values < .05 are denoted in bold. ROI number corresponds to the order of each region in the Seitzman atlas. Coordinates are reported in MNI space.
